# Analyzing Brain Morphology in Alzheimer’s Disease Using Discriminative and Generative Spiral Networks

**DOI:** 10.1101/2021.04.15.440008

**Authors:** Emanuel A. Azcona, Pierre Besson, Yunan Wu, Ajay S. Kurani, S. Kathleen Bandt, Todd B. Parrish, Aggelos K. Katsaggelos, for the Alzheimer’s Disease Neuroimaging Initiative

## Abstract

Several patterns of atrophy have been identified and strongly related to Alzheimer’s disease (AD) pathology and its progression. Morphological changes in brain *shape* have been identified up to ten years before clinical diagnoses of AD, making its early detection more relevant. We propose novel geometric deep learning frameworks for the analysis of brain shape in the context of neurodegeneration caused by AD. Our deep neural networks learn low-dimensional shape descriptors of multiple neuroanatomical structures, instead of handcrafted features for each structure. A discriminative network using spiral convolution on 3D meshes is constructed for the *in-vivo* binary classification of AD from healthy controls (HCs) using a fast and efficient “spiral” convolution operator on 3D triangular mesh surfaces of human brain subcortical structures extracted from T1-weighted magnetic resonance imaging (MRI). Our network architecture consists of modular learning blocks using residual connections to improve overall classifier performance.

In this work: (1) a discriminative network is used to analyze the efficacy of disease classification using input data from multiple brain structures and compared to using a single hemisphere or a single structure. It also outperforms prior work using spectral graph convolution on the same the same tasks, as well as alternative methods that operate on intermediate point cloud representations of 3D shapes. (2) Additionally, visual interpretations for regions on the surface of brain structures that are associated to true positive AD predictions are generated and fall in accordance with the current reports on the structural localization of pathological changes associated to AD. (3) A conditional generative network is also implemented to analyze the effects of phenotypic priors given to the model (i.e. AD diagnosis) in generating subcortical structures. The generated surface meshes by our model indicate learned morphological differences in the presence of AD that agrees with the current literature on patterns of atrophy associated to the disease. In particular, our inference results demonstrate an overall reduction in subcortical mesh volume and surface area in the presence of AD, especially in the hippocampus. The low-dimensional shape descriptors obtained by our generative model are also evaluated in our discriminative baseline comparisons versus our discriminative network and the alternative shape-based approaches.

## 1. Introduction

Advances in magnetic resonance imaging (MRI) have enabled a plethora of non-invasive shape analysis tools and techniques for modeling the human anatomy in high detail, specifically neuroanatomical shape modeling (Ng et al., 2014). Methodological insights in human brain shape analyses have demonstrated powerful utility for their visualization capabilities and valued characterizations of neuropathology and neurodevelopment. Shape-based descriptors have proven to be effective for a variety of tasks such as: segmentation, observing and identifying shape asymmetries, and surface analyses using triangular meshes, each demonstrated by Brignell et al. (2010). Morphological patterns of change in brain structures have often been predictive of different neurodevelopmental and neurodegenerative diseases, such as: schizophrenia, epilepsy (Kim et al., 2013), Lewy bodies, and Alzheimer’s disease (AD) (Shakeri et al., 2016). Neuroanatomical changes in structural MRI have been identified up to ten years before clinical diagnoses in AD (Tondelli et al., 2012). Wachinger et al. (2015) employ BrainPrint to yield extensive characterizations of brain anatomy using structure-specific shape descriptors with samples from the Alzheimer’s Disease Neuroimaging Initiative (ADNI) dataset (Petersen et al., 2010) to identify unique individuals (3000 subjects) with a 99.8% accuracy. Gutiérrez-Becker et al. (2021) demonstrate a strong performance (0.80/0.79/0.78 performance/recall/F1-score respectively) using BrainPrint to classify scans belonging to subjects with AD apart from healthy controls (HCs), which they outperform in a baseline comparison with their own shape descriptors (0.83/0.84/0.82 precision/recall/F1-score respectively) learned on point cloud representations of neuroanatomical shapes.

Working with geometric shape descriptors offers a more robust representation of brain morphology, rather than direct image intensities. The inferences drawn from utilizing shape descriptors are able to remain robust w.r.t intensity changes that may be caused by differing scanner hardware/protocols. A recent development in deep learning (DL), PointNet (Qi et al., 2017a), introduces artificial neural network (NN) architectures designed for operating on 3D point clouds for tasks such as object identification. Gutiérrez-Becker et al. (2021) utilize the point cloud operations from PointNet (Qi et al., 2017a) to construct deep NNs that are trained for AD vs. HC classification on unordered 3D point cloud representations of subcortical brain structures. Their framework is also evaluated on the mild cognitive impairment (MCI) vs. HC classification task, which yields a significant drop in classifier performance due to the high variability within the MCI class, since the detection of MCI is more symptomatic and it is sub-divided into different stages (typically early MCI and late MCI).

Generalizations of successful convolutional neural network (CNN) models to non-Euclidean data types, such as point clouds and triangulated meshes, fall under the wide umbrella of *geometric deep learning* (Bronstein et al., 2017). Similar to 3D voxels (Wu et al., 2016), point clouds (Achlioptas et al., 2018) are intermediate representations of 3D shapes, unlike direct surface representations such as meshes. Despite their high success, voxel-based DL approaches typically suffer from high computational complexity, and point cloud approaches suffer from an absence of smoothness of the data representation (Bouritsas et al., 2019). Polygon meshes are direct and effective surface representations of object *shape*, when compared to voxels. Geometric learning on meshes has only recently been explored (Kolotouros et al., 2019; Litany et al., 2018; Ranjan et al., 2018; Wang et al., 2018; Wickramasinghe et al., 2020) for shape completion, non-linear facial morphable model generation and classification, 3D surface segmentation, and reconstruction from 2D/3D images. A novel representation learning and generative DL framework using spiral convolution on fixed topology meshes, was established with Neural3DMM by Bouritsas et al. (2019) and later improved upon with SpiralNet++ by Gong et al. (2019).

Given the relevance and valued characterizations of brain shape in neuropathology and neurodevelopment, as well as the added value of successful DL methods for shape-driven tasks on 3D point clouds (Qi et al., 2017a,b), we improve upon the work by Gutiérrez-Becker et al. (2021), which operates on unordered point clouds of 3D brain structures. We extend their discriminative networks by working with spiral convolution operators on triangular meshes instead. Similar to Gutiérrez-Becker et al. (2021), we use a *conditional* generative network framework to introduce non-imaging data, particularly AD diagnosis, to analyze the learned morphological patterns of generated meshes w.r.t. diagnostic priors.

Our framework is based upon the spiral convolution operators defined in SpiralNet++ (Gong et al., 2019) and the residual NN framework for image recognition established by He et al. (2016). We quantitatively evaluate the performance of our model in AD/MCI binary classification with an ablation study using different subcortical structure inputs (all structures, per-hemisphere, and per-structure) to analyze the efficacy of incorporating input data from multiple brain regions. Furthermore, we perform a baseline comparison with our spiral framework’s performance with our prior work (Azcona et al., 2020) using spectral graph convolution (Defferrard et al., 2016), and the point cloud approach by Gutiérrez-Becker et al. (2021) on the same AD/MCI classification tasks. Using a conditional variational autoencoder (CVAE) (Sohn et al., 2015) framework, our generative model is used to extract low-dimensional brain shape descriptors that are then used for the same AD/MCI classification tasks. We also experiment with the learned effects of conditioning our generative model on AD diagnosis during training and mesh generation (synthesis).

An interpretation of classifier *reasoning* is often a desired quality of DL frameworks that is often neglected but highly needed, especially in medical image analyses for widespread acceptance or trust. This paper is an extension of our preliminary work (Azcona et al., 2020) where spectral graph convolutional networks (GCNs) (Kipf and Welling, 2017) were used for binary AD classification and we adapted Grad-CAM (Selvaraju et al., 2017) on triangular meshes to provide visually interpretable heatmaps that localize areas on meshes which drive true positive (TP) AD predictions. Given Grad-CAM’s modularity to work with any CNN model, we apply a mesh adaptation of Grad-CAM (Azcona et al., 2020) on the discriminative network in this study.

In summary our contributions are as follows:

1. **A joint framework for improved *in-vivo* disease classification using multiple subcortical structures in a single scan**. A holistic view of brain subcortical anatomy is provided to demonstrate stronger discriminative performance with multiple brain structures. For AD in particular (Frisoni et al., 2010; Klöppel et al., 2008), correspondences across multiple structures are often more robust than studying one organ in isolation, especially in neuroimaging where segmenting multiple subcortical regions is possible from a single MRI volume. AD has also been identified start in localized brain regions (good for early detection) and progressively spreads to multiple brain regions (good for robust detection).
2. **Discriminative spiral networks for improved AD classification on meshes versus prior spectral method**. We demonstrate an improvement in accuracy, precision, recall, and F1-score upon our prior work (Azcona et al., 2020) by using spiral convolution on brain surface meshes for AD classification. Our discriminative spiral network also outperforms alternative shape-based descriptor approaches which operate on intermediate shape representations such as point clouds.
3. **Mesh Grad-CAM adaptation to provide visual reasoning in localized regions of interest (ROIs) on mesh manifolds that drive TP predictions in AD classification**. Our prior adaptation of Grad-CAM (Azcona et al., 2020) was successful in localizing ROIs on meshes for TP predictions from our GCNs. Although a different convolution operator is used in this proposed framework, learned feature maps are still attainable from convolutional layers for generating class activation maps (CAMs) onto input mesh surfaces. These CAMs are a visual interpretation of regions on the along the surface of subcortical structures whose shape is indicative of TP AD predictions by our spiral networks. Our obtained CAMs draw direct correspondences with brain shape deformations tightly correlated with AD pathology.
4. **Conditional generative spiral networks for low-dimensional representation learning on brain mesh manifolds with diagnostic priors**. Our generative CVAE models are able to learn low-dimensional discriminative representations of mesh inputs, which are then evaluated against our proposed discriminative network and prior baseline methods. The morphological effects of conditioning on AD are also analyzed and supported by multiple reports on the neuroanatomical changes in AD progression.

## 2. Related Work

### 2.1. BrainPrint

The shape descriptors in BrainPrint are used in multiple tasks including: (1) subject identification, (2) age and sex classification, (3) lateral asymmetry in brain shape, (4) and potential genetic influences on brain morphology (such as twin analysis) (Wachinger et al., 2015). Using FreeSurfer (Dale et al., 1999; Dale and Sereno, 1993; Fischl et al., 1999a,b, 2002), subcortical labels are used to segment subcortical nuclei (i.e. caudate and hippocampi) from whole brain MRI. Then those subject-specific segmentations are used to create individual triangular meshes of each anatomical structure’s surface. A shape descriptor, referred to as shapeDNA (Reuter et al., 2006), computed from the intrinsic geometry of an object by calculating the Laplace-Beltrami spectrum (Niethammer et al., 2007), is used to compactly represent each structure’s mesh, per scan. Finding shape descriptors that quantify and characterize brain shape are often needed for classification or regression tasks that are dependent on brain shape.

### 2.2. PointNet on 3D neuroanatomical surfaces

DL networks for the shape analysis of neuroanatomical structures using point clouds are introduced by Gutiérrez-Becker et al. (2021) as an improvement on BrainPrint regarding AD pathology. These types of DL approaches naturally scale and benefit in the analysis of large datasets, with potential to learn characteristic variations in large populations. Point clouds are a lightweight representation of 3D surfaces that avoid topological constraints of shapes and are trivial to obtain given a segmented surface. Although computationally robust, their method still operates on and outputs *intermediate representations* of brain shape.

Methods that generate intermediate representations of 3D surfaces (i.e. pixels), are left insensitive to the topological constraints of 3D objects. The output quality of postprocessing steps taken to generate 3D surfaces, like triangular meshes, therefore become dependent on the output quality of the intermediate representations (Marton et al., 2009). In this work, we improve upon the framework established by Gutiérrez-Becker et al. (2021) by working with spiral convolution operators that operate directly on 3D morphable triangular mesh surfaces (Bouritsas et al., 2019) that are registered to a common template topology. We also improve upon their framework by way of residual connections (He et al., 2016) within our classifier, and demonstrate an improvement in classification performance using residual connections within each alternative approach in our baseline classifier comparison.

Additionally, Gutiérrez-Becker et al. (2021) demonstrate a powerful framework for fixed-size point cloud reconstruction and generation using a PointNet CVAE architecture. Although point cloud methods can be compact and robust, they can still lack an underlying smooth structure whose approximation is dependent on the quality of the cloud, whereas surface meshes are still more realistic, less sensitive to noise, and are capable of preserving high-quality 3D geometry generation. In this work, we construct CVAEs using fixed-size surface meshes that are registered to a common template during preprocessing.

### 2.3. Spectral Graph Convolution (ChebyNets)

Morphable meshes (Bouritsas et al., 2019), specifically triangular meshes (TaubinÝ, 2000), are direct surface representations of 3D volumes that can be used for 3D visualization, describing 3D texture, and contextualizing *shape*. By construction, triangular meshes are undirected graphs, with analogous edges, and their intersections are interpreted as vertices. Several studies (Bessadok and Rekik, 2018; Fornito et al., 2015; Göktaş et al., 2020; Gurbuz and Rekik, 2020; Nebli and Rekik, 2020; Yang et al., 2020) have demonstrated that graphs derived from different types of brain-related connectivity, functional or structural, are more robust in accuracy and computation time, versus traditional neuroimaging methods.

Modeling convolution on 3D meshes can be more memory efficient and allow for the processing of higher resolution 3D structures compared to volumetric approaches using 3D CNNs. Our prior work (Azcona et al., 2020) demonstrates an improvement in AD classification with spectral GCNs known as ChebyNets (Defferrard et al., 2016), using a dataset of 3D surface meshes extracted from a subset of T1-weighted MRIs in the subject population used by Punjabi et al. (2019), a volumetric approach. ChebyNets are also implemented by Ranjan et al. (2018) for a generative framework using convolutional mesh autoencoders (CoMA), for generating 3D human faces.

Spectral filtering on graphs (Defferrard et al., 2016; Kipf and Welling, 2017) can come with a number of caveats. Spectral filters are inherently *isotropic* since they particularly rely on the *Laplacian operator*, which performs weighted averages of neighboring vertices:

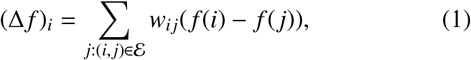

given a shared feature, *f*, on vertices *i* and *j*, and the scalar edge weight, *w*_*i j*_, corresponding to edge *e*_*i j*_ ∈ *ε*, connecting *i* and *j*. Gong et al. (2019) point out that the isotropic nature of spectral filters for undirected graphs is a side effect of needing to design a permutation-invariant operator with a small number of parameters, under the *absence* of a canonical ordering.

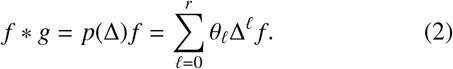

While a “necessary evil” for certain graph learning applications (Bouritsas et al., 2019), spectral graph filters are still basis-dependent and can be rather weak on meshes since they are locally rotational-invariant. On the other hand, spiral convolutional filters take advantage of the fact that meshes are locally Euclidean and a canonical ordering of neighbors for each vertex can be established, such as a spiral sequence starting at an arbitrary vertex. By design, spiral filters are *anisotropic* and have proven to generalize functions on 3D meshes better than spectral methods (Bouritsas et al., 2019; Gong et al., 2019). In our analysis, an ablation study demonstrates an improvement upon AD classification performance with spiral filters, in comparison to the spectral filters defined by Defferrard et al. (2016); originally used in our preliminary work (Azcona et al., 2020).

### 2.4. Generative networks on brain graphs

Several studies have recently investigated using geometric deep learning (Bronstein et al., 2017) for synthesizing brain-related graphs (Bessadok et al., 2019, 2021; Sserwadda and Rekik, 2021; Zhang et al., 2020) using generative adversarial network (GAN) (Goodfellow et al., 2014) inspired frameworks. Other types of generative networks, namely autoencoder-based architectures, have also demonstrated success for neuroimaging applications, such as the work of Choi et al. (2018), where generative models are developed using chronological age and apoE4 genetic traits as conditional features for synthesizing PET scan in relation to AD. In their study variational autoen-coders (VAEs) (Kingma and Welling, 2014) are used to demonstrate a significant effect on apoE4 genetic information in predicting age-related metabolic changes in synthesized PET scans that are then compared to ground-truth follow-up scans.

Autoencoders are neural networks trained to minimize the reconstruct error between their inputs and outputs, separated by *encoder* and *decoder* halves. Traditionally, autoencoders have been used for unsupervised dimensionality reduction or feature learning, since their objective functions for training are typically designed to minimize the reconstructions of its inputs (i.e. mean absolute error).

*Variational* autoencoders (VAEs) (Kingma and Welling, 2014), similarly aim to reconstruct inputs, while also attempting to constrain the latent space of the encoder output to an assumed underlying probabilistic distribution (such as a multivariate Gaussian). Using this assumption, the total objective function used to train VAEs minimize a reconstruction loss term and a latent space regularization term, typically the Kull-back–Leibler (KL) divergence (Joyce, 2011), as a measure of the disparity between the embedding and assumed prior distribution 𝒩 (**0, I**). Once trained, VAEs are valuable in their utility as a generative framework, where new samples can be synthesized by sampling from the assumed prior distribution. CoMA (Ranjan et al., 2018) is built upon a VAE framework for meshes, using spectral GCNs. Their results demonstrate remarkable performance in synthesizing a diversity of facial expressions on 3D morphable meshes, all registered to a common template topology.

As a generative framework, one drawback to VAEs is the lack of control in *targeted* data generation. This can be problematic for tasks dependent on generating specific *types* of samples. *Conditional* variational autoencoders (CVAEs) (Sohn et al., 2015) offer more control by combining variational inference from VAEs with additional conditional priors, w.r.t. each sample, using a simple concatenation step prior to decoding. Based on CoMA and the success of point cloud generation for neuroanatomical shapes (Gutiérrez-Becker et al., 2021), a CVAE framework composed of spiral convolutional learning blocks is used in this study to generate 3D mesh surfaces of neuroanatomical structures by conditioning on AD diagnosis.

## 3. Methods

Data used in the preparation of this article were obtained from the Alzheimer’s Disease Neuroimaging Initiative (ADNI) database (adni.loni.usc.edu). The ADNI was launched in 2003 as a public-private partnership, led by Principal Investigator Michael W. Weiner, MD. The primary goal of ADNI has been to test whether serial magnetic resonance imaging (MRI), positron emission tomography (PET), other biological markers, and clinical and neuropsychological assessment can be combined to measure the progression of mild cognitive impairment (MCI) and early Alzheimer’s disease (AD).

### 3.1. Mesh notation

The input domain of our data is represented using triangular mesh manifolds, ℳ= (𝒱, ε, ℱ), for the corresponding finite set of vertices, edges, and faces for each mesh. In graph signal processing (Wu et al., 2020b), meshes are treated as undirected graphs, where a feature vector of *F* features at vertex *i* is defined by a row vector **x**_*i*_ ∈ ℝ^*F*^, for *N* vertices. To encapsulate all of the shared features on the vertices of a single mesh, we use the vertex feature matrix, **X** ∈ ℝ^*N*×*F*^.

The shared features on the vertices of the input meshes to our models are the corresponding *x, y, z* coordinates (3 features total) of each vertex in the corresponding subject’s native 3D space. The meshes used in our study are all registered to a common mesh template of the subcortical structures utilized in our prior work (Azcona et al., 2020; Besson et al., 2020; Wu et al., 2020a). The meshes in this study use a shared topology (same number of vertices/edges). However, the *positions* of vertices vary across samples therefore representing a span of different 3D morphology for each sample using the same template of connectivity. An analogous set-up with Euclidean data could be a 3D array of voxels, where the features at each voxel is the corresponding location of the voxel in the subject’s native space.

### 3.2. Mesh extraction

Beginning with the obtained T1-weighted MRIs, FreeSurfer v6.0 (Fischl, 2012) is used to denoise each scan, then followed by B1 field homogeneity corrections and intensity/spatial normalization. Seven subcortical structures per hemisphere were segmented (amygdala, nucleus accumbens, caudate, hippocampus, pallidum, putamen, thalamus) and modeled into surfaces using SPHARM-PDM.

Next, surfaces were inflated, parameterized to a sphere, and registered to a common spherical surface template using a rigid-body registration to preserve the subcortical (Besson et al., 2014, 2020) anatomy. Then, surface templates were converted to triangular meshes following a triangulation scheme. A scalar edge weight, *w*_*i j*_, was assigned to each edge, *e*_*i j*_, connecting vertices *i* and *j*, using their geodesic distance, *ψ*_*i j*_, along the surface s.t.

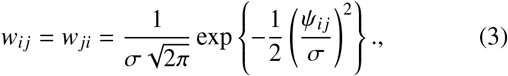

for σ = 2 selected ad-hoc.

As done in our prior work (Azcona et al., 2020), surface templates were parcellated using a hierarchical bipartite partitioning of their corresponding mesh. Beginning with the initial mesh representations of densely triangulated surfaces, we used spectral clustering to define two partitions. These two partitions were then each separated, yielding four child partitions, and repeated so forth. This process was repeated until the average distance across neighboring partitions was 2.5mm. Given one partition, we define the central vertex of a partition as the vertex whose centrality was the highest. The distance across two partitions was defined as the geodesic distance (in mm) across the central vertices of each partition. Two partitions are considered neighbors if at least one vertex in each partition were connected.

Finally, partitions were numbered so that partitions 2*p* and 2*p* + 1 at level *L*, had the same parent partition *p* at level *L* − 1. Therefore, for each level a mesh was obtained s.t. the vertices of the mesh were the central vertices of the partitions and the edges across neighboring vertices were weighted following Eq.3 This serves as an improvement upon the work of Defferrard et al. (2016) to ensure that no singleton is ever produced by mesh coarsening operations for the subcortical structures. At the finest level, a single mesh sample had a total of *N*_0_ = 14, 848 vertices to represent all the subcortical structures (see Table 1 for vertex counts per structure per hemisphere).

**Table 1:** Number of vertices per subcortical structure per hemisphere.

| Structure | # Vertices, $N$ |
| --- | --- |
| amygdala | 512 |
| caudate | 1,024 |
| hippocampus | 2,048 |
| nucleus accumbens | 256 |
| pallidum | 512 |
| putamen | 1,024 |
| thalamus | 2,048 |

### 3.3. Spiral sequences on triangular meshes

Next, we provide an illustrated clarification of spiral sequences on 3D morphable brain meshes (Figure 1), which are at the core of the learning framework introduced by Gong et al. (2019). Given an arbitrary triangular mesh and an arbitrarily-selected vertex we call the *center vertex*, a spiral sequence can be naturally enumerated by following a spiral pattern around the center vertex. First, a spiral orientation is fixed (clock-wise or counter-clockwise) and a random starting direction is selected from the center vertex. Following the convention of Gong et al. (2019), orientations for all spiral generations were fixed to *counter-clockwise* and an *arbitrary* starting direction w.r.t. each vertex was used.

**Figure 1:**
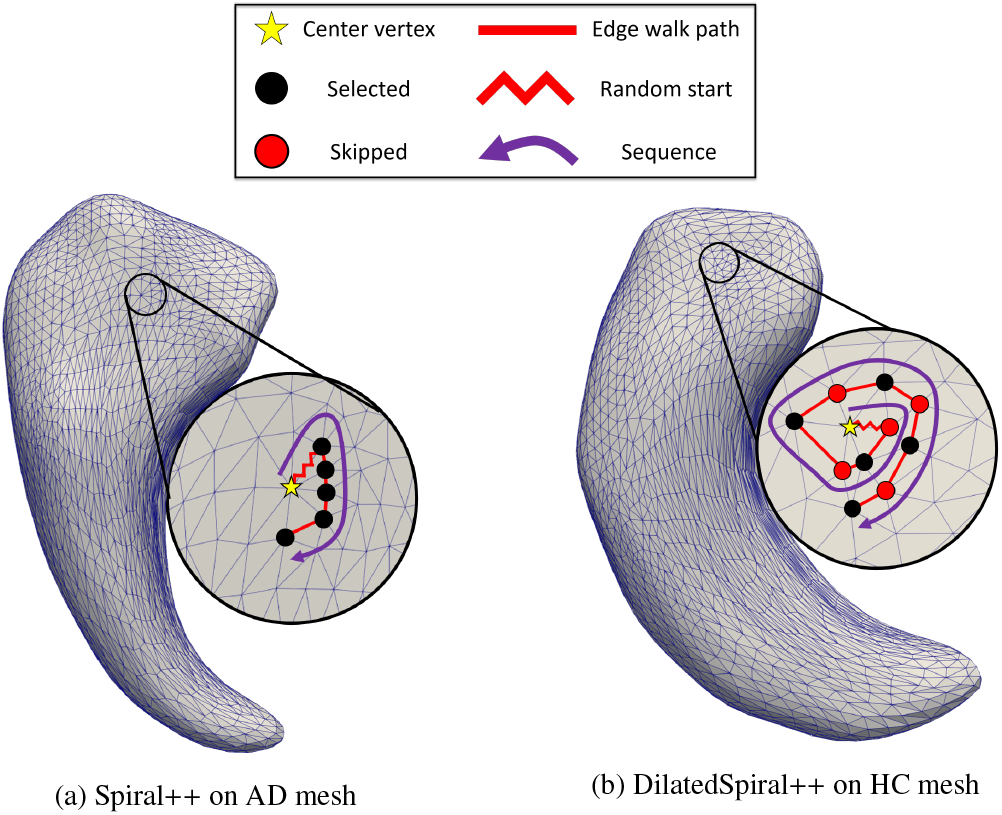
Examples of spiral sequences established on left hippocampi triangular meshes from randomly selected scans of a subject with Alzheimer’s disease (left) and a healthy control (right). Note that in using dilation, the receptive field of the kernel supports exponential expansion without increasing the support-size/length of the spiral kernel (Gong et al., 2019). In each example, a spiral sequence of 6 selected vertices are generated including the center vertex.

Specifically, a *k*-ring and a *k*-disk around a center vertex *v* is defined as:

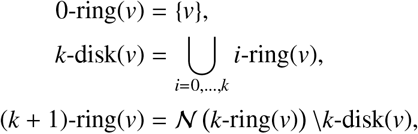

where 𝒩(*V*) is the set of all vertices adjacent to any vertex in set *V*. A spiral sequence of length *ℓ* at vertex *v* is defined as *S* (*v,ℓ*); a *canonically ordered* set of *ℓ* vertices from a concatenation of *k*-rings. Only part of the last ring is concatenated in this definition, in order to ensure a fixed-length serialization. Formally, the spiral sequence is defined as:

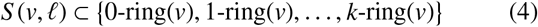

The spiral sequences defined in SpiralNet++ (Gong et al., 2019) show remarkable advantages to a high-level feature representation for each vertex in a consistent and robust way when spirals are *frozen* during training. By this we mean that spiral sequences are generated *only once* for each vertex on the template mesh, instead of randomly generated sequences for every vertex per epoch, as was done by Lim et al. (2018). Since the 3D mesh samples used in this study are all registered to a common template topology, the same spiral sequences can be used for every sample. By design, this automatically generates the topology of the convolutional filter on each vertex of the template mesh, analogous to the assumed rectangular topology of convolutional filters with standard 2D Euclidean CNNs.

### 3.4. Spiral convolution

Convolutional neural networks (CNNs) applied on 2D/3D images defined on standard Euclidean grids (Deng et al., 2009; LeCun et al., 1989) are designed using 2D/3D rectangular convolutional kernels that slide across the images and map *F*_*in*_ input feature maps to *F*_*out*_ output feature maps. An extension of this application on data types in irregular domains such as graphs, is typically expressed using neighborhood aggregation (Corso et al., 2020; Xie et al., 2020) or message passing (Gilmer et al., 2017) schemes.

Using the convention defined in Section 3.1, with 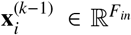 denoting the feature vector of *F*_*in*_ features at vertex *i* and 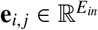 denoting the (optional) *E*_*in*_ features on edge *e*_*i,j*_ connecting vertex *i* to vertex *j* at layer (*k* −1), message passing neural networks are typically defined s.t.

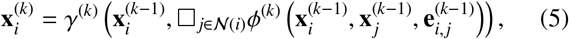

where □ represents a differentiable permutation-invariant operation (i.e. sum, mean or max), and *γ*^(*k*)^ and *ϕ*^(*k*)^ denote differentiable kernel functions such as Multi-Layer Perceptrons (MLPs) (Fey and Lenssen, 2019). CNNs defined for data types that exist in standard Euclidean grids have a clear one-to-one mapping. However for data types in irregular domains such as graphs, correspondences are defined using neighborhood connectivity for each vertex and weight matrices dependent on the kernel functions, *γ* and *ϕ* at each layer.

Using the spiral sequence serialization discussed in Section 3.3, we can define convolution on meshes in an equivalent *canonical* manner to Euclidean CNNs that is *anisotropic* by design. Following the convention of Gong et al. (2019), the spiral convolution operator is defined as

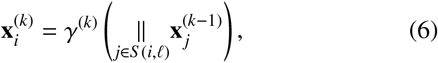

where *γ* denotes MLPs and ∥ is the concatenation operation applied on the shared features of the vertices of spiral sequence *S* (*i, ℓ*) centered at vertex *i*.

The *dilated* extension (Yu and Koltun, 2016) of spiral convolution using the dilated spiral sequence (depicted in Figure 1) can also be applied to meshes by uniformly sub-sampling during spiral generation, with the preprocessing parameter *d* denoting a uniform sampling of every *d* − 1 vertices along the spiral sequence until *ℓ* vertices are selected.

### 3.5. Mesh sampling (down/up-sampling)

Traditional Euclidean CNNs, typically use a hierarchical multiscale learning structure, typically employed for learning global and local context, using a combination of convolutional and pooling/up-sampling layers. To mimic this behavior, we use mesh sampling/coarsening operators (Ranjan et al., 2018) that define analogous down-sampling and up-sampling of mesh vertices within a neural network.

As mentioned in Section 3.1, vertex feature matrices for meshes with *N* vertices and *F* shared features, are denoted **X** ∈ ℝ^*N*×*F*^. The 3D mesh samples in this work use *F* = 3 input dimensionality, however convolutions applied on mesh features within the neural network can result in features with different dimensionality. Therefore, in this section we use *F* to generalize our definition.

The in-network down-sampling of a mesh, with *N* vertices, is performed using the down-sampling matrix, **D**∈ {0, 1} ^*M*×*N*^, and up-sampling with **U** ∈ ℝ^*N*×*M*^, for *N* > *M*. The down-sampling matrix, a sparse matrix, is obtained by contracting vertex pairs iteratively that maintain surface error approximations using quadric matrices (Garland and Heckbert, 1997). The vertices of the down-sampled mesh are essentially a subset of the original mesh’s vertices, 𝒱_*d*_ ⊂ 𝒱. Each element of **D**(*p q*) ∈ {0, 1} denotes whether the *q*-th vertex is kept during down-sampling, with **D**(*p, q*) = 1, otherwise discarded with **D**(*p, q*) = 0, ∀ *p*.

To remain loss-less, the up-sampling operator is built during the generation of the down-sampling operator. Vertices retained during down-sampling are kept for up-sampling s.t. **U**(*q, p*) = 1 iff **D**(*p, q*) = 1. Vertices *q*∈𝒱 that are discarded during down-sampling, for **D**(*p, q*) = 0, ∀*p*, are mapped into the down-sampled triangular mesh surface by using barycentric coordinates. This is specifically done by projecting *q* into the closest triangle (of vertices *i, j*, and *k*) of the down-sampled mesh surface, denoted by 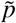, and determining the barycentric coordinates, 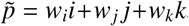, where *i, j, k* ∈ 𝒱 and *w*_*i*_ +*w*_*j*_ +*w*_*k*_ = 1. Using these weights, we update **U** s.t.

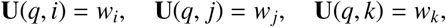

otherwise, **U**(*q, l*) = 0.

The features on the vertices retained from a down-sampling operation for the new mesh are obtained via sparse matrix multiplication in

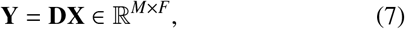

for **X** ∈ ℝ^*N*×*F*^. In a synonymous way, the vertices on the output mesh of an up-sampling operation are obtained as an inverse operation to down-sampling via the sparse matrix multiplication

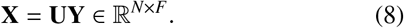

### 3.6. Spiral brain mesh networks

#### 3.6.1. Residual learning blocks (ResBlocks)

Motivated by the success of residual deep learning frameworks (He et al., 2016) for image recognition, the NN models used in this work are based on a residual learning architecture composed of “residual learning blocks” (ResBlocks) depicted in Figure 2. These function by adding the output from a previous block to the output of the current block. This methodology was demonstrated to allow for the training of deeper NN architectures, with the intuition that adding more layers allows for progressively learning more complex features within the architecture (He et al., 2016; Szegedy et al., 2016).

**Figure 2:**
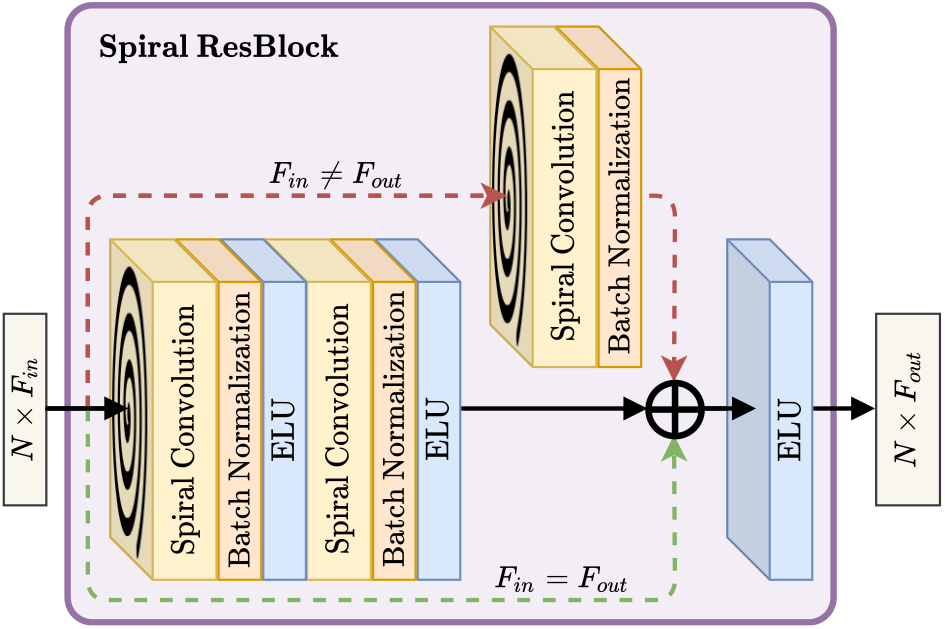
Residual learning block (ResBlock) module used in this SpiralNet++ inspired architecture. Batch normalization (depicted in orange) is applied after spiral convolution (depicted in yellow). The top (red) branch of the ResBlock uses spiral convolution followed by batch normalization as an identity linear mapping tool to map the *F*_*in*_ features of the input vertices to the *F*_*out*_ features acquired by the main branch. Otherwise, the input of the ResBlock is added to the main branch output (green). An element-wise ELU(·) function is used within the hidden layers and as the final activation of the ResBlock.

A spiral convolutional layer maps *F*_*in*_ ⟼ *F*_*out*_ features for every vertex in the input mesh using MLPs applied on the spiral sequence, *S* (*i, ℓ*) of each vertex, *i*. Analogous to traditional convolution with padding to preserve the size of input feature maps, spiral convolution on meshes also preserves dimensionality since *S* (*i, ℓ*) is defined for every input vertex. Therefore, the number of vertices, *N*, is still preserved in the output vertex feature matrix, 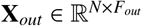.

A frequent problem in DL with training deep NNs is the *internal covariate shift* in the distribution of inputs to layers (Ioffe and Szegedy, 2015) within a model. Batch normalization (BN) is used after each spiral convolution operation within our Res-Blocks as a way to prevent our networks from “forever chasing a moving target,” by standardizing the inputs to layers within the network. This follows the convention used of other successful DL architectures related to computer vision (He et al., 2016; Szegedy et al., 2016).

An important hyperparameter for training deep networks is the choice of activation function for the hidden layers and output layer. He et al. (2016), used the rectified linear unit (ReLU), defined as

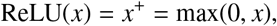

for their residual learning framework. DL architectures using ReLU, are prone to suffering from the common “dying ReLU” problem where hidden layer outputs heavily saturate to zero (Lu et al., 2020), leading to zero-valued gradients, making learning more difficult. We circumvent this by using the exponential linear unit (ELU) activation function (Clevert et al., 2015) defined as

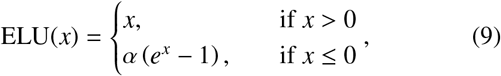

for α = 1 in this work.

#### 3.6.2. Convolutional mesh encoder

Using ResBlocks introduced in Section 3.6.1 and mesh down-sampling, described in Section 3.5, we introduce the convolutional *encoder* module used by our discriminative and generative spiral networks, illustrated by Figure 3. As illustrated, input feature matrices are embedded to 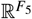 latent vectors using the encoder defined as the sequential stack: {ResBlock(*ℓ*_1_, *d*_1_, *F*_1_) → MS(↓ 2) → ResBlock(*ℓ*_2_, *d*_2_, *F*_2_) → MS(↓ 2) → … → ResBlock(*ℓ* _5_, *d*_5_, *F*_5_) → MS(↓ 2) → GAP_*N*_}, where

**Figure 3:**
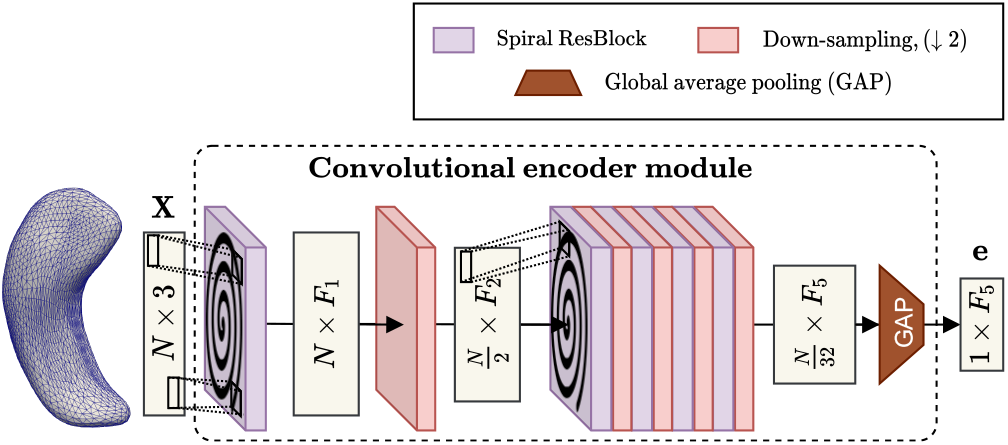
Convolutional mesh encoder module made up of a sequential stack of alternating spiral convolution and down-sampling layers (5 each). The *i*-th ResBlock maps *F*_*i*_ features onto the vertices of the respective input. Each down-sampling layer coarsens the input vertex count down by a factor of 2. After the final down-sampling layer, global-average pooling (GAP) is applied over the vertex dimension to reduce the output embedding down to 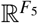.

- *ℓ*_*r*_, *d*_*r*_, *F*_*r*_ are the spiral lengths, dilation, and number of filters for all convolutional layers w.r.t. the *r*-th ResBlock,
- MS(↓ 2) is shorthand for “mesh-sampling down by a factor of 2” (down-sampling), and GAP_*N*_ is the global-average pooling operation (Selvaraju et al., 2017).

On meshes, GAP_*N*_ is essentially just an averaging operation over the node dimension, as depicted in Figure 3.

Note that since the input mesh is down-sampled 5 times within the module, each time by a factor of 2, the number of vertices after the final down-sampling operation is 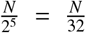. This module is used as the first step for both our discriminative and generative networks, described in Sections 3.6.4 and 3.6.5 respectively.

#### 3.6.3. Convolutional mesh decoder

The convolutional mesh *decoder* module, depicted in Figure 4, applies a synonymous backwards transformation of the encoder module described in Section 3.6.2. Following Figure 4 and starting with an arbitrary vector **z** ∈ ℝ^*k*^, first a fully-connected (FC) layer maps 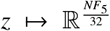. This output is then reshaped to get a feature matrix in 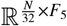, representing the *F*_5_ features on the 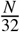 vertices at the coarsest level of our meshes. The rest of the decoder module is defined as the sequential stack: MS(↑ 2) → ResBlock(*ℓ*_5_, *d*_5_, *F*_5_) → MS(↑ 2) → ResBlock(*ℓ*_4_, *d*_4_, *F*_4_) →… → MS(↑2) ResBlock(*ℓ*_1_, *d*_1_, *F*_1_) → SpiralConv(*ℓ*_1_, *d*_1_, 3). Here *ℓ*_*r*_, *d*_*r*_, and *F*_*r*_ are the *same* corresponding values used in the encoder module. An additional spiral convolutional layer (SpiralConv) with 3 filters is used at the end (with no activation) to obtain the reconstruction, 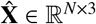, with 3 features per vertex (corresponding *x, y, z* coordinates). This module is only utilized within the generative network described in Section 3.6.5, where the task is to output 3D meshes.

**Figure 4:**
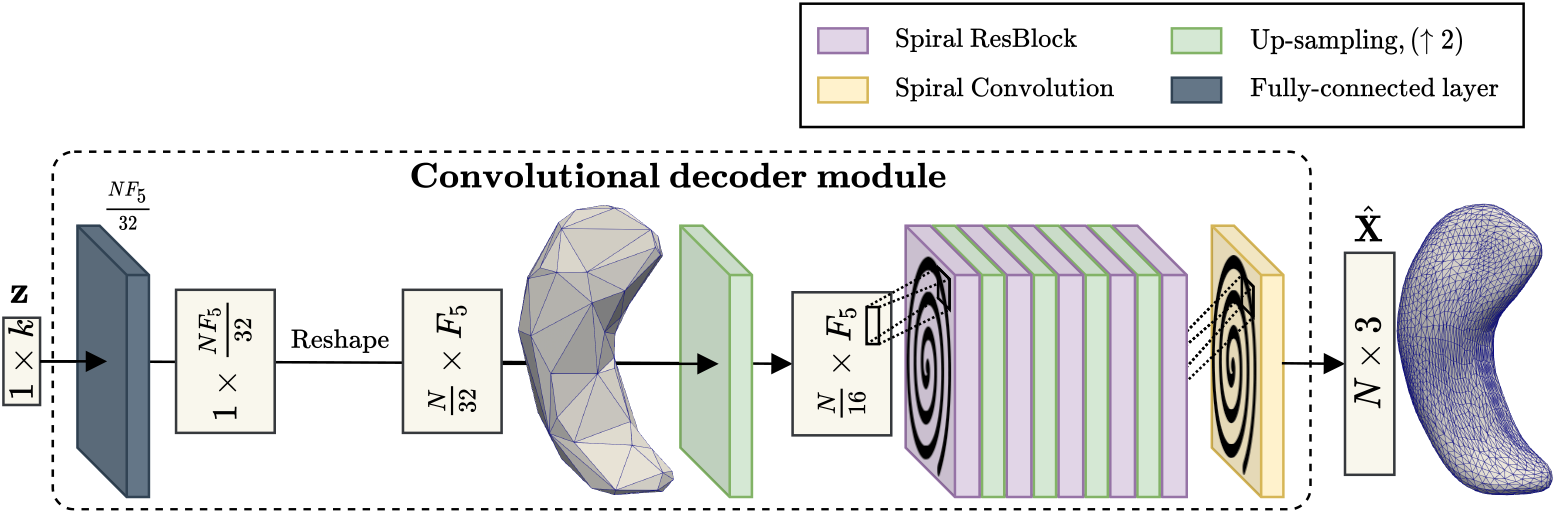
The mesh decoder module first uses a FC layer and reshaping to map the input vector, **z** ∈ ℝ^*k*^ to a feature matrix for meshes at the coarsest level in 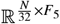. Alternating up-sampling and ResBlock layers (5 each) are used after. An additional spiral convolutional layer with 3 filters and no activation function is used to project the penultimate *N* × *F*_1_ feature matrix back to *N* × 3 for the respective 3D mesh reconstruction.

#### 3.6.4. Discriminative network

Following the point cloud discriminative network convention established by Gutiérrez-Becker et al. (2021), we construct our discriminative networks using the encoder module (Figure 3) in series with a MLP that uses BN and a ELU activation functions after each FC layer, as depicted by Figure 5. The goal of this network is to learn mesh features given an input feature matrix, **X** ∈ R^*N*×3^, and a spiral convolutional operator that exploits the locally-Euclidean topology of 3D mesh manifolds. These learned mesh features are then global-average pooled and used within a MLP for predicting the target variable, *y*.

**Figure 5:**
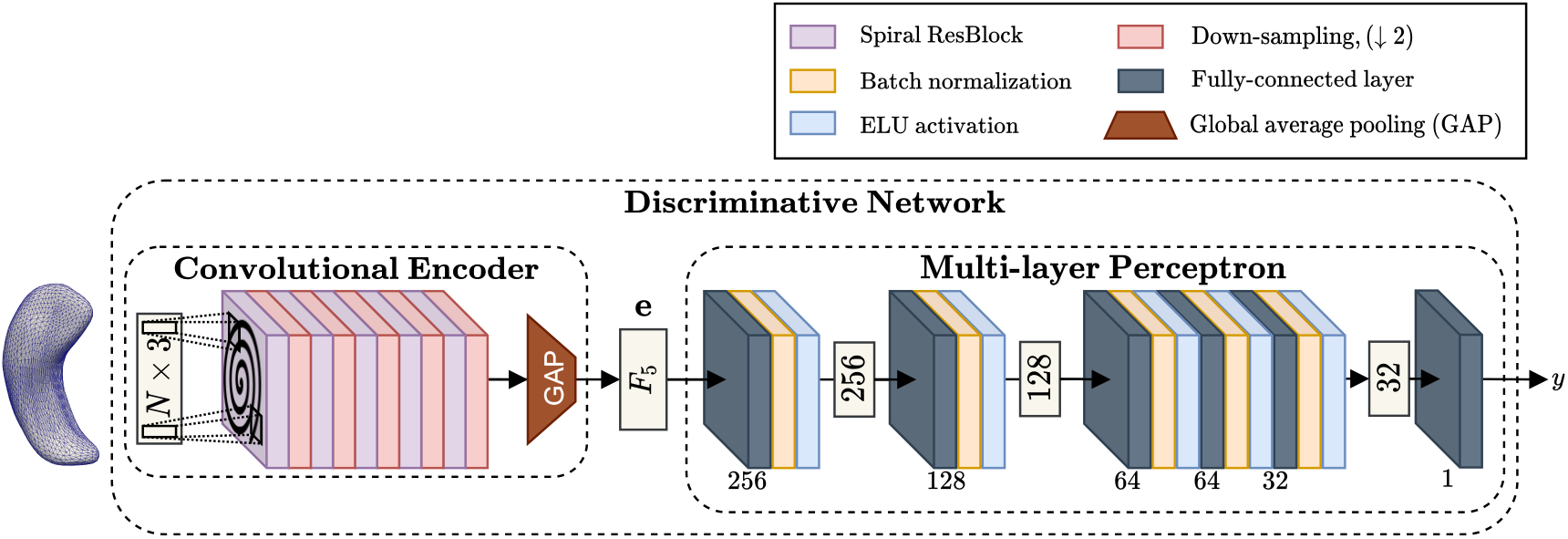
End-to-end discriminative spiral network given a 3D mesh input with feature matrix **X** ∈ ℝ^*N*×3^. Batch normalization is used after each MLP layer, followed by an ELU(·) activation. Given the output of the convolutional encoder, 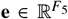, the MLP predicts the target, *y*, from the embedding for a particular sample. For binary classification, we apply a sigmoid function after the final layer to output a probability for each sample.

In this work, we use the discriminative network for binary classification, therefore we apply a sigmoid function, 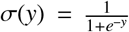, on the predicted targets to get the probability of a positive label given the corresponding 3D mesh manifold. Traditionally, for binary classification tasks such as disease prediction, the positive binary label, (1), pertaining to the pathology, is typically the positive class in opposition to the healthy control label, (0). Our discriminative network can be trained in an end-to-end supervised manner by optimizing a standard binary cross-entropy (BCE) loss

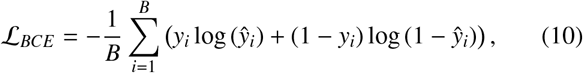

where *y*_*i*_ and *ŷ*_*i*_ are the ground-truth labels and predicted probabilities (output of sigmoid) respectively, for a given sample, *i*, in a batch of *B* samples.

#### 3.6.5. Generative network (CVAE)

Based on the CoMA architecture by Ranjan et al. (2018), our CVAE model uses a convolutional decoder on mesh samples that share a topology at different hierarchical levels of coarsening, described in Section 3.6.3. Following Figure 6, first a convolutional encoder, *E*, (Section 3.6.2) is used to compress input samples, **X** ∈ ℝ^*N*×3^, down to a latent vector, 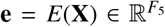. Next, **e** is mapped to a “mean vector,” µ ∈ ℝ^*k*^ and a “standard deviation vector,” σ∈ ℝ^*k*^, using two parallel fully-connected (FC) layers. These vector outputs are then used for variational inference during training with the “reparameterization trick” for VAEs (Kingma and Welling, 2014). Taking 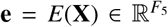, where *e*_*i*_ ∈ **e**, we vary each component of the latent vector as *z*_*i*_ = µ_*i*_ + ϵσ_*i*_ ∈ ℝ^*k*^, where ϵ ∼ 𝒩 (0, 1), therefore assuming a multivariate Gaussian distribution, *Q* (**z** |[**z**]**X**), that can be sampled from.

**Figure 6:**
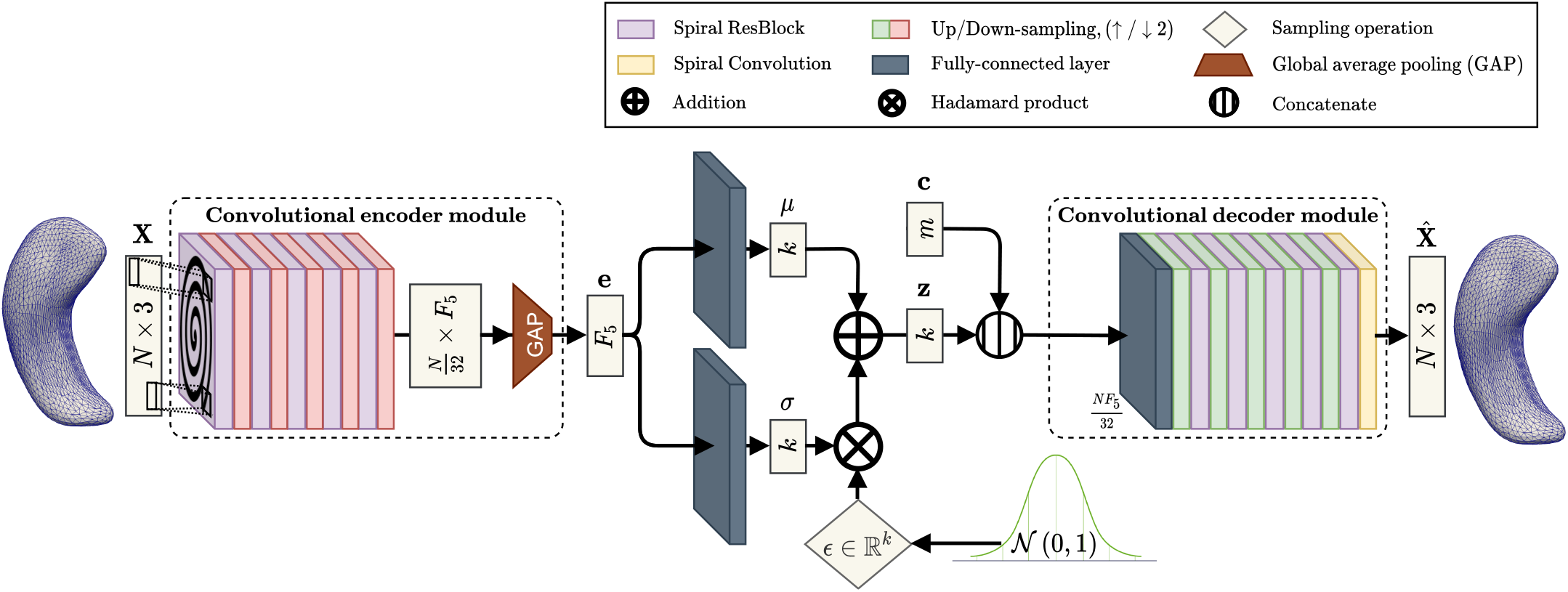
End-to-end generative model based on spiral convolutional CVAE architecture. During inference, a mesh sample, **X** ∈ ℝ^*N*×3^, is first encoded to 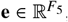, using the encoder, *E*. This encoding is then used to sample, **z** ∈ ℝ^*k*^, from the prior distribution, *Q* (**z**|**X**), assumed to be a multivariate Gaussian. Next **z** is concatenated with the conditional vector, **c** ∈ ℝ^*m*^, and a reconstruction is generated using the decoder 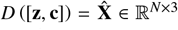. During generation, we sample from 𝒩 (0, 1) for each varied component of **z**, concatenate the sample with a given conditional **c**, and start at the decoder to generate a new sample, *D* ([**z, c**]).

Next we concatenate a random sample, **z**, with the associated conditional vector **c**, to generate the mesh reconstruction, 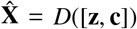. As done by Ranjan et al. (2018) for CoMA, our spiral CVAE is trained by minimizing the loss

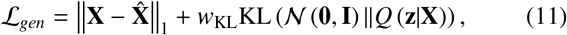

with w_KL_ = 0.001, selected ad-hoc, acting as a weight on the KL divergence loss. The first term (reconstruction) minimizes the MAE between the reconstruction and ground truth sample, and the second term (KL divergence) acts as regularizer on the latent space by adding the constraint of a unit Gaussian prior with zero-mean on the latent space distribution, *Q*(**z** |**X**).

Once trained, synthesizing new samples is simple. Since the KL divergence constrains the latent space to a unit Gaussian, we generate new samples with our decoder by sampling a ℝ^*k*^ vector from the unit Gaussian prior and concatenating it with a conditional prior vector, **c** ∈ ℝ^*m*^, as a “specification mechanism” on the type of sample we want to synthesize.

### 3.7. Grad-CAM mesh adaptation

In our preliminary work (Azcona et al., 2020), we adapt a visualization tool known as Grad-CAM (Selvaraju et al., 2017) to provide an interpretable localized heatmap, that weighs the “importance” of areas in an image that are indicative of certain predictions after a model is trained. In our prior work, class activation maps (CAMs) were extracted from our discriminative model to highlight areas, directly onto surfaces, that led to true positive (TP) predictions in AD binary classification. Wu et al. (2020a) also use this mesh adaptation of Grad-CAM to highlight the areas of the cortex and subcortical structures that were most indicative for predicting fluid intelligence in children and adults.

Following the convention of Selvaraju et al. (2017), first the gradients of the class scores (logits prior to softmax or sigmoid) w.r.t. to the feature maps at the last convolutional layer (prior to GAP) are extracted. Using these gradients, GAP is applied on each feature map, per-class, to extract the “neuron importance weights,” 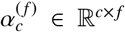, whose formulation was readapted for meshes s.t.

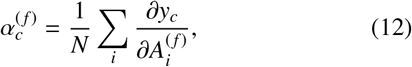

where *y*_*c*_ corresponds to the class score of class *c*, and 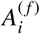 to the value at vertex *i* in feature map *A*^(*f*)^ ∈ℝ^*N*^. The set of neuron importance weights, 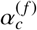, is then projected back onto each feature map, *A*^(*f*)^, to compute the CAMs, *M*_*c*_ for each class, *c*, s.t.

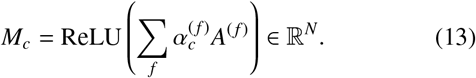

ReLU is applied to the linear combination of maps because we are only interested in the features that have a positive influence on the class of interest (Selvaraju et al., 2017).

As a consequence of multiple down-sampling operations within our discriminative network’s architecture, extracted CAMs w.r.t. the number of vertices at the final convolutional layer are smoother and less focused to specific surface locations depending on the number of layers. Therefore, they are up-sampled back to the same number of vertices as the input, using a trivial interpolation, for a direct “overlay” onto the original input mesh.

## 4. Experiments

We evaluate our discriminative and generative spiral networks for several supervised and unsupervised tasks respectively. First, we introduce the 3D structural neuroimaging dataset and describe our convention for assigning the appropriate *in-vivo* diagnosis labels for each mesh sample (Section 4.1.1). Next we detail the preprocessing parameters used within our experiments for generating the spiral sequences at each level of mesh coarsening (Section 4.1.2).

In Section 4.2, we conduct an experiment with our discriminative model to analyze the efficacy of incorporating input data from multiple subcortical structures for binary AD/MCI classification. Our results demonstrate a clear advantage to the joint modeling of multiple subcortical structures, as opposed to using a single hemisphere or single structure. In Section 4.2.2, we provide a baseline comparison to alternative shape-based operators, in place of spiral convolution, for the same binary classification tasks. In Section 4.2.3, CAMs are generated for samples that are correctly classified as AD by our spiral discriminative network. These CAMs fall in accordance with previous reports of morphological changes observed in the brain correlated with AD. Our CAMs support our classification results by producing visual transparency into our discriminative network’s reasoning for true positive AD classification.

Lastly, in Section 4.3, we evaluate the effect of conditioning on AD diagnosis for our generative models w.r.t. each subcortical structure. Our generative network’s results demonstrate that our model captures morphological differences in the presence of AD for some of the subcortical structures, particularly the hippocampi and amygdala nuclei, which are in accordance with previous autopsy reports that highlight patterns of atrophy associated to AD.

### 4.1. Dataset and pre-processing

#### 4.1.1. ADNI dataset

In this study, we use 8,665 T1-weighted 3D MRI volumes from the Alzheimer’s Disease Neuroimaging Initiative (ADNI) dataset, corresponding to 1,266 unique subjects. For each scan, we associate the healthy control (HC), mild cognitive impairment (MCI), or Alzheimer’s disease (AD) labels given up to 2 months after the corresponding scan in ADNI. This is done as a precaution to ensure that each diagnosis had clinical justification. Our dataset consists of 2,758/3,959/1,948 samples for the HC/MCI/AD labels respectively.

Each discriminative model in this work is designed to classify pathological (AD/MCI) scans apart from HCs. To ensure that scans from the same subject do not appear in different sets, all data splits (train/test/validation) in this study, shuffle samples by *subject* identifiers instead of *scan* identifiers. We randomly split our data into training/testing sets (85%, 15%) across subjects, and use a 5-fold cross-validation across the subjects within the training set in our analyses.

Meshes are extracted from each T1-weighted MRI sample using the mesh extraction preprocessing method described in Section 3.2. Each subcortical region is represented using an independent surface mesh with the corresponding number of vertices described in Table 1, per hemisphere. Using the trimesh (Dawson-Haggerty et al., 2019) library in the Python (Van Rossum and Drake Jr, 1995) programming language, a mesh object for one hemisphere of a subcortical region is represented using a vertex feature matrix, **X**∈ R^*N*×3^, described in Section 3.1, and a corresponding set of faces, ℱ, which is a set of 3-element tuples where each tuple indexing the vertices that make up the corresponding triangular face on the mesh.

The vertex feature matrix of a mesh sample containing a single bilateral subcortical region (i.e. LH/RH hippocampi) is constructed using a row-wise concatenation (vertical stacking) of the vertex feature matrix for each hemisphere of the corresponding subcortical region. The sets of faces for each hemisphere of the same subcortical region are merged to create one set faces for the bilateral subcortical region sample type. For mesh samples representing a single hemisphere or all subcortical regions, the corresponding vertex feature matrix and set of faces is obtained using the same vertical stacking and merging process. Each mesh sample type can be interpreted as an undirected graph with described by vertex feature matrix, **X**, and the corresponding set of faces, ℱ. Therefore, 14, 848 vertices are used to represent a single mesh sample for all subcortical regions, 7, 424 vertices for a single hemisphere, and 2*N* vertices per subcortical region, for each corresponding *N* in Table 1.

#### 4.1.2. Spiral sequence and mesh-sampling generation

Following the encoder module described in Section 3.6.2 and depicted in Figure 3, the topology of spiral sequences at each level of mesh coarsening is only preprocessed *once*. In-order, the spiral lengths, *ℓ*_*r*_, used for the spiral filters within the *r*-th ResBlock of the encoder are 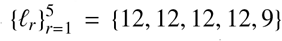, with the corresponding dilation parameters, 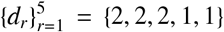. These parameters are used in reverse-order for the ResBlocks within the the convolutional decoder, depicted in Figure 4.

Following the steps in Section 3.5, down/up-sampling matrices were generated *once* to represent surfaces in this study at multiple hierarchical levels while still preserving context at each level. Again following the structure of the encoder (Figure 3), we specifically up/down-sample meshes within the architecture by a factor of 2 for each mesh sampling operation. At each level of coarsening, spiral sequences are generated once using the template mesh.

### 4.2. Discriminative model predictions

Discriminative models and hyperparameter tuning were evaluated using the 5-fold cross-validation on the training set, as explained in Section 4.1.1, for two separate experiments. In our first experiment, we conduct an experiment with our discriminative model to analyze the efficacy of incorporating input data from multiple subcortical structures for binary AD/MCI classification, in comparison to input data from a single hemisphere or single structure. In our second experiment, we analyze the performance of alternative *shape*-based classifiers in comparison to our proposed method. We report the results on the test set for each classification task. The number of filters, per convolutional layer, at the *r*-th ResBlock, within the encoder is 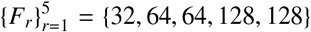. A binary cross-entropy (BCE) objective function was used to train all discriminative models using the AdamW (Loshchilov and Hutter, 2017) optimizer with a learning rate of 2 × 10^−4^, learning rate decay of 0.99 for every step, and a batch size of 16 samples per batch over 200 epochs. In addition to the BCE loss, the weights of the network were also *L*_2_-regularized with a weight decay of 1 × 10^−5^.

#### 4.2.1. Subcortical structural ablation study

First, we perform binary classification tasks on different combinations of subcortical structures to classify scans using the diagnostic labels provided by ADNI. The first task is to classify HC scans apart from those belonging to subjects with AD, meanwhile the second task looks at HC vs. MCI classification. For each task, we use the same discriminative spiral network (architecture and number of parameters) from Section 3.6.4, and train each model on the same task, each with a varied combination of input structures. Classifiers are trained and compared with: (a) single-structure (both hemispheres), (b) single-hemisphere, and (c) all-structure mesh inputs for each sample.

Table 2a summarizes the results of the experiments on **Alzheimer’s Disease (AD) Classification** across variations of subcortical structure inputs. The discriminative model’s performance gradually improves with the inclusion of more subcortical regions. In particular, an improvement in classifier performance is observed when an entire hemisphere (an input with multiple subcortical regions), is used versus using both hemispheres of a single subcortical region. The discriminative model performs best when *all* subcortical regions (the largest input option) are used as input. The discriminative model trained on the left hemisphere (LH) slightly outperforms the model trained on the (RH) in both Area Under the Curve (AUC) statistics, which may also be indicative of the way AD pathology is typically diagnosed. The left hemisphere of the human brain is tightly associated to language function (i.e. grammar, vocabulary, and literal meaning) (Corballis, 2014), which is often used as a metric for the clinical diagnosis of AD.

**Table 2:** Binary classification results using same discriminative model. Precision, recall, and F1-score are reported w.r.t. a classification threshold of 0.5. For a global measure over different thresholds we also report the Area Under the Receiver Operating Characteristic Curve (ROC-AUC) and the Area Under the Precision-Recall Curve (PR-AUC) for each case.

| Structure | Threshold = 0.5 |  |  | AUC |  |
| --- | --- | --- | --- | --- | --- |
|  | Precision | Recall | F1 | ROC-AUC | PR-AUC |
| all structures | **0.877** | 0.834 | **0.855** | **0.906** | **0.895** |
| left hemisphere | 0.827 | 0.700 | 0.758 | 0.893 | 0.874 |
| right hemisphere | 0.737 | 0.798 | 0.766 | 0.887 | 0.863 |
| amygdala | 0.788 | **0.850** | 0.818 | 0.900 | 0.891 |
| caudate | 0.524 | 0.655 | 0.582 | 0.699 | 0.592 |
| hippocampus | 0.682 | 0.722 | 0.702 | 0.812 | 0.708 |
| nucleus accumbens | 0.610 | 0.674 | 0.640 | 0.774 | 0.690 |
| pallidum | 0.543 | 0.644 | 0.589 | 0.700 | 0.556 |
| putamen | 0.642 | 0.637 | 0.639 | 0.780 | 0.705 |
| thalamus | 0.611 | 0.723 | 0.662 | 0.780 | 0.703 |

(b) Healthy control (HC) vs. Mild cognitive impairment (MCI)
| Structure | Threshold = 0.5 |  |  | AUC |  |
| --- | --- | --- | --- | --- | --- |
|  | Precision | Recall | F1 | ROC-AUC | PR-AUC |
| all structures | 0.613 | **0.712** | 0.659 | 0.612 | **0.693** |
| left hemisphere | 0.629 | 0.616 | 0.622 | 0.589 | 0.649 |
| right hemisphere | 0.645 | 0.631 | 0.637 | **0.622** | 0.691 |
| amygdala | 0.643 | 0.561 | 0.599 | 0.607 | 0.689 |
| caudate | 0.628 | 0.565 | 0.595 | 0.578 | 0.635 |
| hippocampus | 0.597 | 0.705 | 0.647 | 0.549 | 0.622 |
| nucleus accumbens | 0.573 | 0.625 | 0.598 | 0.503 | 0.597 |
| pallidum | 0.597 | 0.698 | 0.643 | 0.533 | 0.617 |
| putamen | 0.602 | 0.551 | 0.575 | 0.529 | 0.618 |
| thalamus | **0.646** | 0.677 | **0.661** | 0.617 | 0.593 |

AD follows a different trajectory than normal aging (Nelson et al., 2011). Language and memory problems like forgetfulness can be correlated with normal aging, however the types of memory problems that occur with AD dementia are more severe and typically begin to interfere with “everyday” activities, which is not a part of normal aging. One example: forgetting where you put your glasses, can be indicative of disorganization, forgetfulness, or normal aging. However, forgetting *what* those glasses are used *for* (their utility) is not a part of normal aging. Like many anomaly detection problems in medical imaging, where it is important to anticipate pathological events that occur less times than the healthy control, *precision-recall* statistics (see Tables 2a and 2b) are often stronger for measuring classification performance when there is a class imbalance and the class of interest belongs to the smaller population.

There exists strong evidence for certain patterns of atrophy for different neuroanatomical structures at different stages of AD progression (Dickerson et al., 2001). Early involvement of the entorhinal cortex, hippocampus, and amygdala in AD progression have been reported consistently in the literature (Braak and Braak, 1991; Dubois et al., 2007; Klein-Koerkamp et al., 2014; Ledig et al., 2018). Our results in Table 2a suggest a stronger performance in AD classification given the *shape* of the amygdala or hippocampus alone compared to the other subcortical structures. Most importantly, these results also demonstrate that a holistic approach incorporating multiple subcortical regions improves AD classification.

Table 2b demonstrates the results of **Mild Cognitive Impairment Classification**. An expected drop in performance occurs for MCI classification, compared to AD. This behavior is expected due to the MCI group’s variability, given its detection being more symptomatic and it is also sub-divided into several stages. Detecting MCI is important because people with MCI are more likely to develop AD than those without. Unlike the fluidity of the MCI pathological spectrum, AD progression is endemic and symptoms worsen with time. However, methods related to neuroplasticity exist to potentially slow/mitigate its progression, making the early detection of AD desirable.

#### 4.2.2. Spiral discriminator baseline comparison

Given the improvement in AD classification using input data from multiple subcortical regions for our discriminative model, we compare our model’s performance with other baseline *shape-based* classifiers on the same dataset. We evaluate four different methods to perform the same discriminative tasks as Section 4.2.1: (1) the discriminative network in this work, (2) the same discriminative module with spectral graph convolution in-place of spiral convolution, (3) the end-to-end discriminative network by Gutiérrez-Becker et al. (2021), and (4) a MLP trained on the latent space features of the generative network in this work.

### Spectral networks set-up

For the spectral convolutional (ChebyNet) (Defferrard et al., 2016) network comparison, we demonstrate an improvement in performance with BN and a residual learning architecture by training and evaluating multiple learning architectures. We construct (1) a ChebyNet using the same architecture as the discriminative network in Figure 5, but with ChebyNet layers, BN, and ELU activations in place of the Spiral ResBlocks, and another network using “ChebyNet ResBlocks,” where spiral convolution operations within a Res-Block are replaced with ChebyNet layers. For a fair comparison, we use the same network depth as the spiral discriminator, the same number of output features per convolutional layer, and a Chebyshev polynomial of degree *K* = 6 for each spectral convolutional layer (Defferrard et al., 2016). The second MLP-half of each ChebyNet model follows the same MLP architecture model follows the same MLP architecture used within our spiral discriminative model (Figure 5).

### Point cloud networks set-up

To utilize the same dataset on this method, we drop the edges of our 3D meshes and treat the surface vertices as point clouds representing the surface/shape of the subcortical structures. The shared MLPs within the architecture of the discriminative model constructed by Gutiérrez-Becker et al. (2021) to operate on point clouds, are identically implemented using 1D convolutional layers with a kernel size of 1 (Qi et al., 2017a,b). For consistency in adopting the PointNet-inspired model for a fair comparison, we use the PointNet layers described by Gutiérrez-Becker et al. (2021), and construct the same discriminative network in Figure 5, with point cloud operations in place of spiral operations. We construct a PointNet discriminator with (1) 1D convolutional layers + no activation following Gutiérrez-Becker et al. (2021), in place of the ResBlocks, (2) the same PointNet model in addition to BN + ELU activations after each convolutional layer, and (3) a final variant with “PointNet ResBlocks” following the same style as the Spiral ResBlocks and “ChebyNet ResBlocks” in the spectral set-up. The second MLP-half of each PointNet model follows the same MLP architecture used within our spiral discriminative model (Figure 5).

### Generative model latent space set-up

A generative model (Section 3.6.5) was constructed with 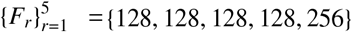 for the corresponding output feature maps of the model’s encoder and decoder Spiral ResBlocks. We found it best to compress mesh samples down to a latent space using ℝ ^16^ components for each subcortical structure, therefore resulting in **z** ∈ ℝ ^112^ for all subcortical structures. A binary one-hot encoding vector is used for the condition vector **c** ∈ ℝ ^2^, w.r.t. the diagnosis label for each sample.

The generative network was trained by optimizing the loss function in Equation 11 and using *L*_2_-regularization, weighted by 1 × 10^−5^, on the network’s parameters. The AdamW (Loshchilov and Hutter, 2017) optimizer is used with a learning rate of 2 × 10^−4^, learning rate decay of 0.99 for every step, and a batch size of 8 samples per batch over 500 epochs of training. Once trained, a MLP following the same architecture as the second MLP in the discriminative network (Figure 5), is trained on the latent space shape descriptors (i.e. **z**) of the corresponding samples, using the same data splits as the other baseline comparisons. This MLP is also trained using the AdamW (Loshchilov and Hutter, 2017) optimizer, with the same training parameters as the rest of the discriminative baseline models.

### AD model comparison

For the AD binary classification task, the model comparison results in Table 3a demonstrates that our spiral discriminative model used in the previous ablation study (Section 4.2.1) outperforms all the baseline models in precision, recall, and F1 score for a 0.5 binary classification threshold. Our model also outperforms the baseline models in both Area Under the Curve (AUC) statistics, particularly the Precision-Recall Curve (PR-AUC) indicating an overall improvement in precision, recall, and F1 score across multiple classifier thresholds in [0, 1].

**Table 3:** Baseline comparison of binary classifiers for HC versus AD/MCI classification.

(a) Healthy control (HC) vs. Alzheimer’s disease (AD)
| Model | Threshold = 0.5 |  |  | AUC |  |
| --- | --- | --- | --- | --- | --- |
|  | Precision | Recall | F1 | ROC-AUC | PR-AUC |
| SpiralResNet (Ours) | **0.877** | **0.834** | **0.855** | **0.906** | **0.895** |
| Generative (Ours) | 0.703 | 0.771 | 0.735 | 0.851 | 0.769 |
| ChebyNet | 0.487 | 0.644 | 0.555 | 0.664 | 0.580 |
| ChebyResNet | 0.740 | 0.757 | 0.748 | 0.869 | 0.837 |
| PointNet | 0.791 | 0.798 | 0.795 | 0.803 | 0.786 |
| PointNet+BN+ELU | 0.802 | 0.776 | 0.789 | 0.798 | 0.774 |
| PointResNet | 0.842 | 0.814 | 0.828 | 0.836 | 0.822 |

(b) Healthy control (HC) vs. Mild cognitive impairment (MCI)
| Model | Threshold = 0.5 |  |  | AUC |  |
| --- | --- | --- | --- | --- | --- |
|  | Precision | Recall | F1 | ROC-AUC | PR-AUC |
| SpiralResNet (Ours) | **0.613** | 0.712 | 0.659 | 0.541 | **0.693** |
| Generative (Ours) | 0.595 | 0.776 | 0.673 | 0.524 | 0.629 |
| ChebyNet | 0.602 | 0.820 | **0.694** | 0.542 | 0.612 |
| ChebyResNet | 0.591 | **0.827** | 0.689 | 0.521 | 0.610 |
| PointNet | 0.590 | 0.789 | 0.676 | 0.528 | 0.615 |
| PointNet+BN+ELU | 0.595 | 0.826 | 0.692 | **0.557** | 0.639 |
| PointResNet | 0.601 | 0.702 | 0.648 | 0.542 | 0.616 |

The spectral classifier without residual connections (ChebyNet in Table 3a) performs the worst overall. However, with the addition of the residual learning framework by using “ChebyNet ResBlocks,” we see an improvement in performance across all metrics for the ChebyResNet model; in fact, it ranks second-highest in both AUC scores behind our spiral model. In our prior work (Azcona et al., 2020), ChebyResNets were used for the same AD binary classification task on the same subcortical structures used in this study, in addition to the corresponding white and pial cortical surface meshes for each sample. In that work, ChebyResNet outperformed the baseline classifiers, demonstrating an improvement in performance by directly learning on surface meshes with spectral graph convolution. In this work, ChebyResNet outperforms the PointNet variants, indicating again an improvement in performance over non-surface mesh approaches.

The bare PointNet model, without activation functions or a residual framework, performed better across all metrics (shown in Table 3a), in comparison to the bare ChebyNet classifier. The PointNet models progressively improves overall with the addition of BN + ELU activations, and with the residual learning framework. The PointResNet model does outperform the ChebyResNet model in precision, recall, and F1-score given a 0.5 binary classification threshold, however not in AUC statistics taken over several thresholds in [0, 1].

### MCI model comparison

Like our structure ablation experiment, we see a drop in performance for all discriminative models in binary MCI classification. The same network set-up used for AD classification was used in this evaluation, treating MCI as the positive label. Our SpiralResNet classifier achieves the highest PR-AUC in MCI classification when compared to the baseline methods. The overall drop in performance for MCI classification for all models in this experiment is the same behavior analyzed in the previous experiment.

#### 4.2.3. Class activation maps for AD classification

Using the pre-trained SpiralResNet classifier trained on all the subcortical structures, we generate class activation maps (CAMs), using our Grad-CAM adaptation on meshes, for each AD sample in the test set that is correctly classified by our model (TP predictions), given a 0.5 classifier threshold. CAMs for the TP samples are then averaged and projected onto the vertices of the subcortical template (Besson et al., 2014, 2020). Mesh faces are colored using an interpolation based off the CAM values at the vertices of each corresponding triangle (each face has 3 corresponding vertices). The color-map scale used to visualize the TP CAM in Figures 7a and 7b highlights areas along the surface by their magnitude of influence, ordered from least to greatest, in binary AD classification with our trained discriminative model.

**Figure 7:**
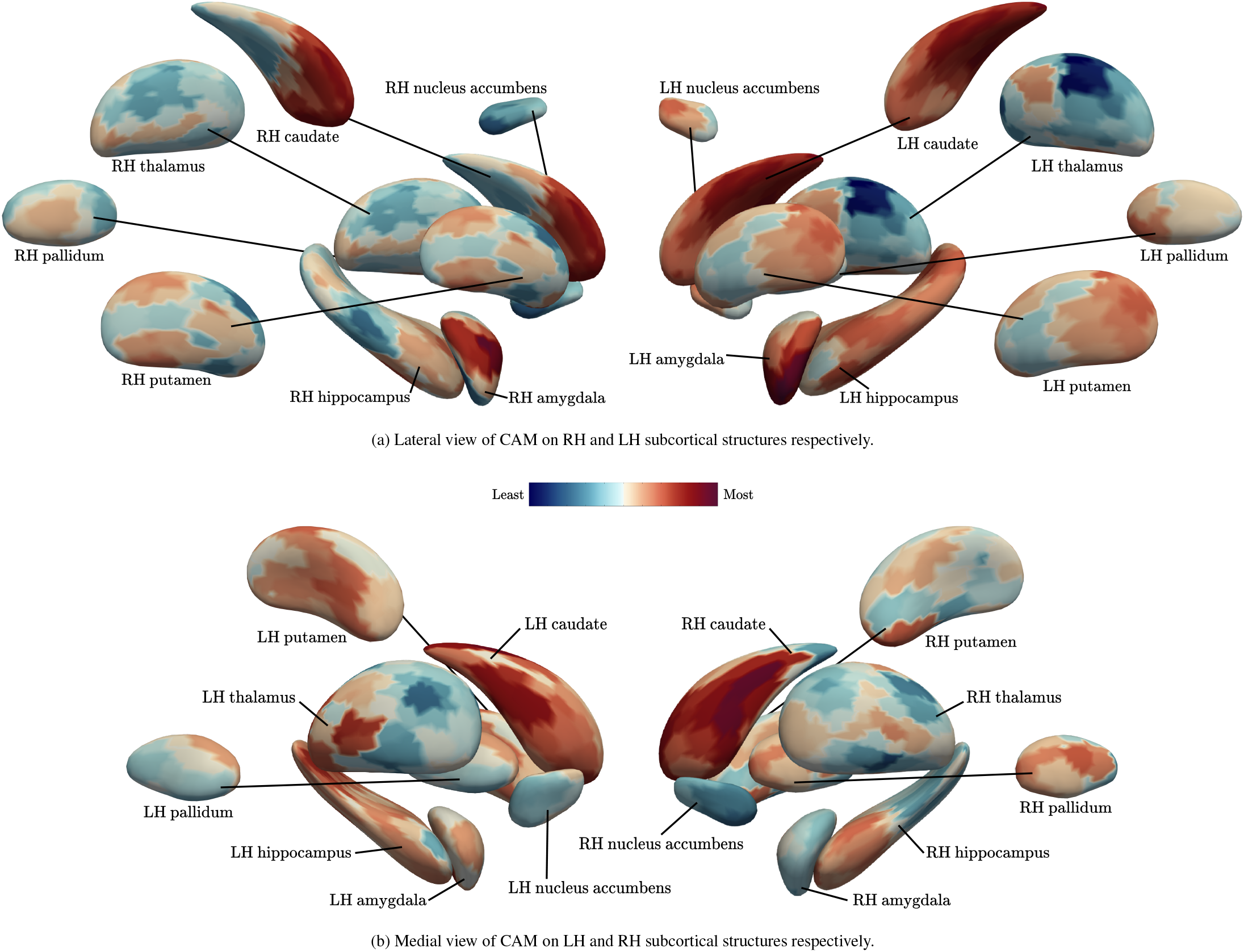
Average of class activation maps (CAM) for true positive predictions by the SpiralResNet discriminative network proposed in this work. A CAM is generated for each TP prediction and their average is projected onto the subcortical structure template mesh by Besson et al. (2014). Provided are lateral (a) and medial (b) views of the CAM projected on the template, which follows the color-scale map which at the center of the two subfigures.

Aligning with our discriminative model’s results in the structure ablation study, we observe a strong involvement of hippocampus and amygdala shape in AD vs. HC classification. In AD, it has been demonstrated that cortical atrophy occurs earlier and progresses faster in the LH than in the RH (Long et al., 2013; Thompson et al., 2007). Wachinger *et al*. demonstrated a significant leftward asymmetry in cortical thinning (mainly in the temporal lobe and superior frontal regions) with an increase in hippocampal asymmetry, which remains consistent with previous findings demonstrating an asymmetric distribution of amyloid-*β* (Frings et al., 2015), a protein in the brain that is thought to be toxic and naturally occurs at abnormal levels in the brains of subjects living with AD.

Both caudate structures are also highlighted as indicative of TP classifications, again with a similar leftward asymmetry. In particular, there is an emphasis on the tail of the left caudate nucleus. This observation falls in line with the findings of (Barber et al., 2002) where both the left and right caudate nucleus were smaller in volume for patients with dementia compared to age-matched healthy controls (HC); in fact, their findings show that the left caudate volume difference was significant in AD subjects (*p* < 0.01). In a recent study looking at shape differences in the ventricles of the brain w.r.t. AD, Ferrarini et al. (2006) show that the areas adjacent to the anterior corpus callosum, the splenium of the corpus callosum, the *amygdala*, the thalamus, the tails of the *caudate* nuclei, and the head of the *left caudate* nucleus are all significantly affected by AD and also highlighted within our generated CAMs.

Volume reductions in the putamen, hippocampus, and thalamus volume were observed by de Jong et al. (2008), adhering to the potential left putamen involvement depicted in Figure 7b. On the left hippocampus structure particularly, we see widespread involvement of the structure with most of the predictive activity occurring at the tail of the left hippocampus and roughly around the CA1 subfield, also reported by Gutiérrez-Becker et al. (2021).

On average, we observe an asymmetry towards the CAMs of the LH structures as more indicative of AD than the RH, even with a trained on both hemispheres at once. Our ablation study also demonstrates an improvement in classifier AUC performance (Table 2a) with using the LH versus the RH in AD classification. Several studies point towards a left lateralization of brain atrophy in AD (Long et al., 2013; Thompson et al., 2007), however Derflinger et al. (2011) argue that brain atrophy in AD is asymmetric rather than lateralized and that data suggesting leftward lateralization may be a result of selection bias. This may be due to the fact that clinical scores used to diagnose AD are primarily language-based, resulting in a potential bias towards a selection of patients already with left-lateralized atrophy (Keilp et al., 1996).

### 4.3. Diagnostic conditioning on generative model

Differences in output generation w.r.t. to AD diagnosis was done with the point cloud generative models by Gutiérrez-Becker et al. (2021). Shape variations their model associates to the presence of AD are measured with point-to-point metrics like *L*_1_ distance. Choi et al. (2018), also experiment with modifying their CVAE’s condition vectors to generate synthetic PET images and forecast future age-related metabolic changes. Predicted regional metabolic changes were correlated with the real changes in their follow-up data. In this work, we observe changes in mesh surface area, *A*, and volume, *V*, w.r.t. to the HC and AD labels, given the same latent vectors for the set of HC samples. The shape descriptors learned by our generative model that are used in our discriminative model evaluation (Table 3a) demonstrate potential in encoding complex shape variations using a low-dimensional embedding.

For our final evaluation, we use the same CVAE architecture used by our generative model in Section 4.2.2 to construct a generative model w.r.t. each subcortical structure, using **z** ∈ ℝ^16^ as the dimension of the latent space for each model. For each CVAE model we use a binary one-hot encoding w.r.t. to the AD versus HC labels in our dataset as the condition vector, **c** ∈ ℝ^2^, to analyze the effect of conditioning on AD diagnosis *per structure*. Each CVAE model is trained following the same training parameters and AdamW optimizer used for training the generative network in our baseline classifier comparison (Section 4.2.2).

First we train each generative network on the entire dataset of HC and AD samples. Next, we extract latent space embedding (i.e. **z** ∈ ℝ^16^ for each subcortical structure) of each HC sample in the dataset. With the latent space shape descriptor of each structure for each HC sample, we analyze the effect of changing the HC label to AD before the decoding step of each generative network to see how diagnosis affects the generated output.

Based on the literature regarding changes in the hippocampus shape as a result of AD, Figure 8 qualitatively depicts some of the hippocampus results in four randomly selected (originally HC) samples. Qualitatively, we observe a “thinning” in hippocampus volume for either hemisphere, particularly shown in the examples of the second (LH) and third (RH) columns in Figure 8. The histograms spread throughout Figures 9-12 quantitatively depict the observed corresponding volumes, *V*, and surface areas, *A*, with using the HC samples and changing the diagnosis during decoding. The volume of a watertight mesh is determined using a surface integral, and the surface area is determined as the sum of the areas of all the triangles on a mesh surface.

**Figure 8:**
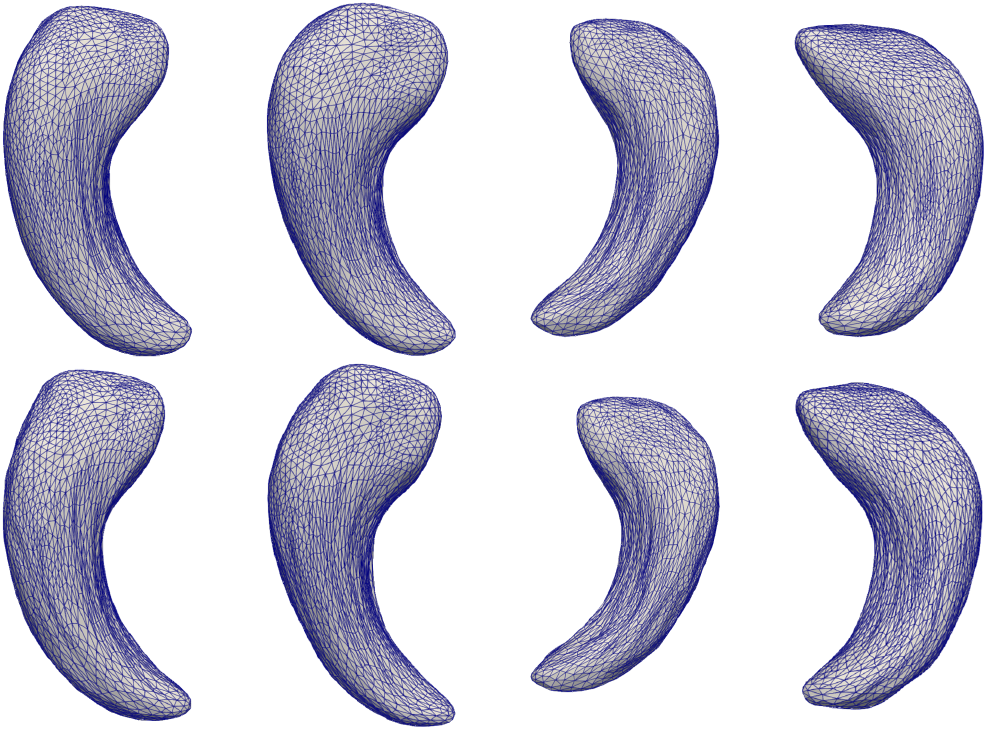
Dorsal views of the left and right hippocampus surfaces generated using proposed generative CVAE model on ADNI dataset. For a given latent space vector, **z**, a 3D mesh is generated by conditioning on the HC (top row) or AD (bottom row) label that is passed along to the decoder along with **z**. Each column corresponds to a different HC sample.

**Figure 9:**
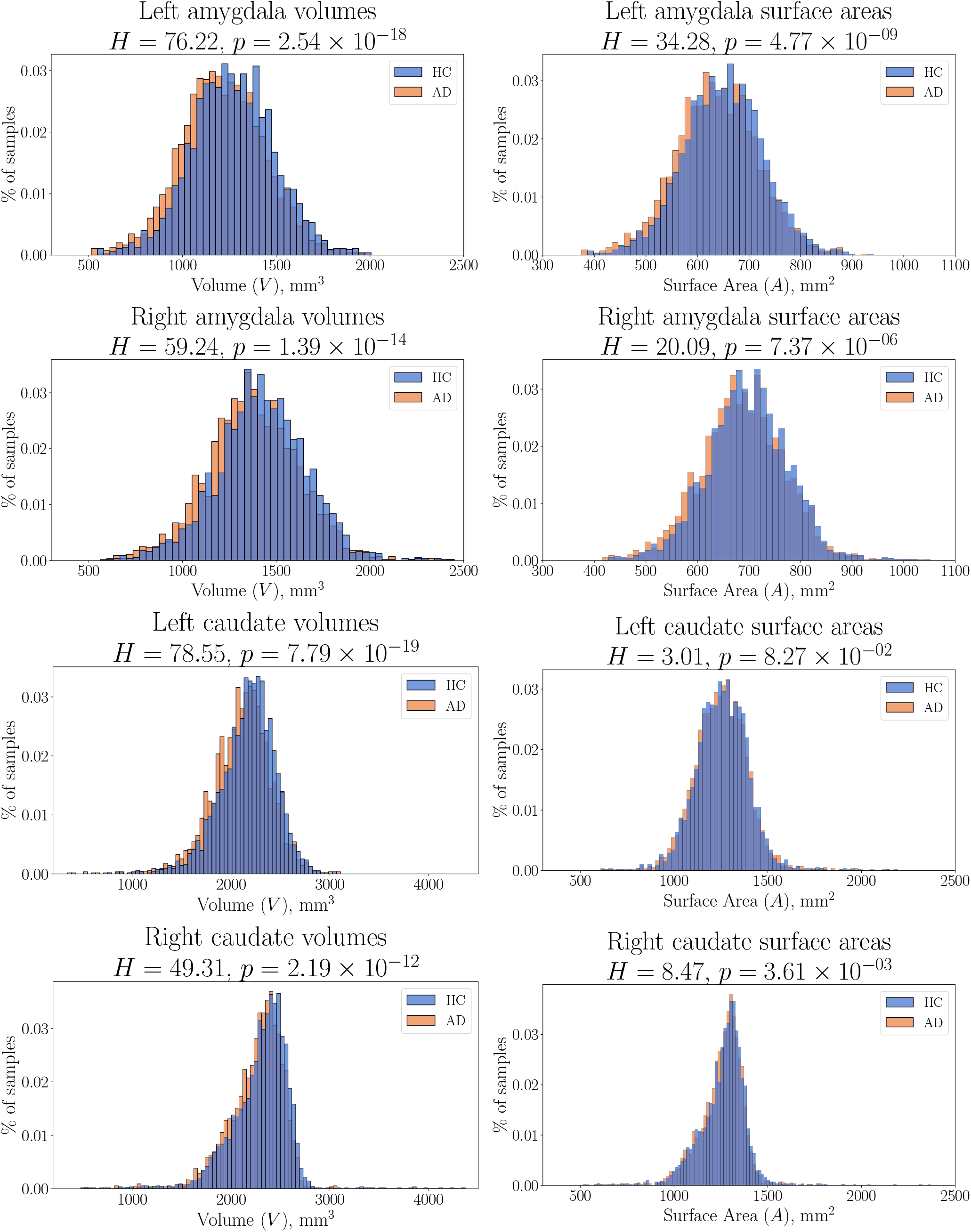
Observed changes in output volume and surface area for amygdala (first two rows) and caudate (bottom two rows).

**Figure 10:**
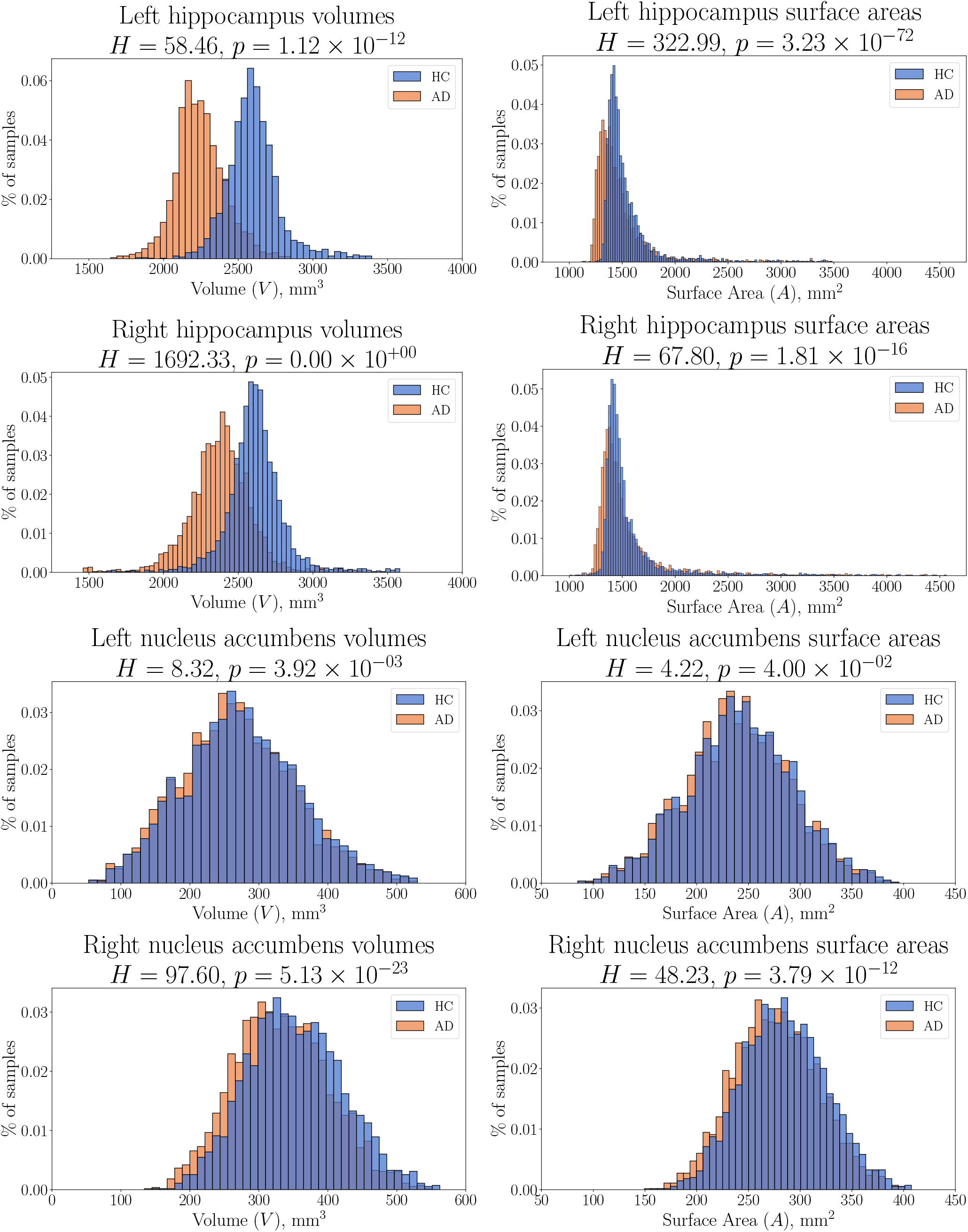
Observed changes in output volume and surface area for hippocampus (first two rows) and nucelus accumbens (bottom two rows).

**Figure 11:**
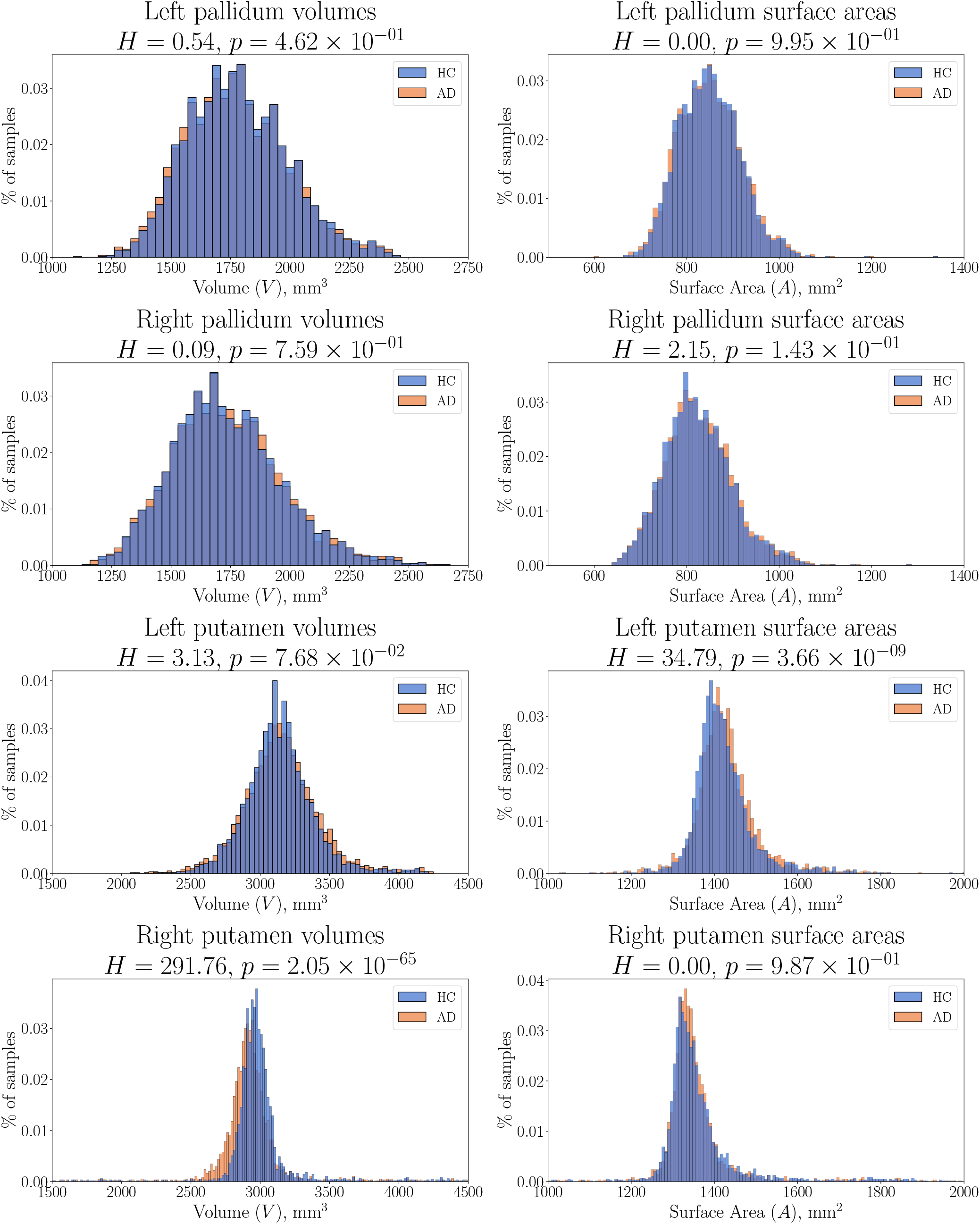
Observed changes in output volume and surface area for pallidum (first two rows) and putamen (bottom two rows).

**Figure 12:**
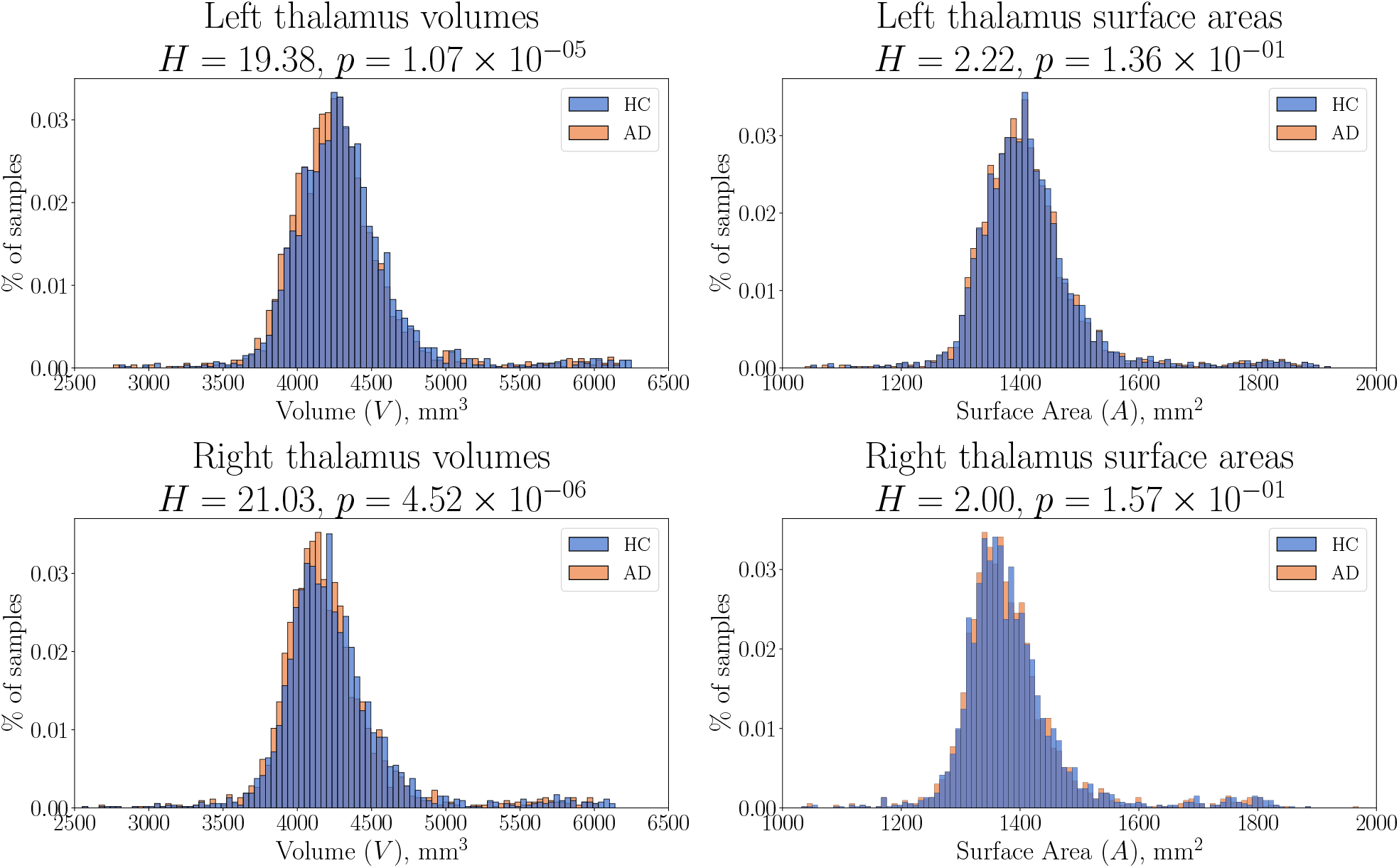
Observed changes in output volume and surface area for thalamus.

Given that the diagnosis labels are categorical and we are analyzing the effect of conditioning the generative shape model using these labels, we use the non-parametric Kruskal-Wallis *H*-test (McKight and Najab, 2010) to measure the statistical significance of differences in the output of the model w.r.t. each label. For each histogram, we report the corresponding *H*-value and *p*-value. For the left putamen, left pallidum, and right pallidum, differences in the volumes of generated outputs are not statistically significant (*p* > 0.05). For the left caudate, left nucleus accumbens, left/right pallidum, right putamen, and left/right thalamus, there is no statistical significance (*p* > 0.05) in the differences in surface area.

For each of the remaining subcortical structures, a reduction in volume and surface area is the most common observation, especially in both hippocampi (*p* ≪ 0.001). We hypothesized our generative model would learn to reduce the hippocampus and amygdala structures, areas that are highly correlated with language, memory, frontal executive function scores. Our results for the remaining structures are in accordance with the expected shrinking of each structure in the presence of AD, coinciding with previous autopsy reports in AD progression (Braak and Braak, 1991; Mann, 1991).

### 4.4. Summary of experiments

Our results for *in-vivo* AD vs. HC classification with spiral networks on brain surface meshes demonstrate the powerful discriminative advantage in learning surface representations of subcortical brain structures. Spiral CNNs are demonstrated to outperform recent methods which operate on point cloud representations or use spectral graph convolution on the same template-registered meshes in this study. To the best of our knowledge, the spiral network method proposed in this study is the only state-of-the-art (SOTA) approach that exploits the locally-Euclidean properties of vertices distributed across a surface to design learnable anistropic filters that improve AD classification w.r.t. subcortical structure *shape*. Our results demonstrate a clear advantage to incorporating multiple subcortical regions, as opposed to input data from a single subcortical region or hemisphere.

The CAMs obtained using our discriminative model draw direct correspondences with the literature regarding localized areas of deformation related to AD pathology. Paired with our discriminative model, our framework combines localized contextual visualization together with classification results. More often, a modular visualization method that provides context to a discriminative model’s predictions *without making architectural changes to the model*, is highly desirable for establishing appropriate trust in predictive models.

Furthermore, the results of our generative model demonstrate the potential for using diagnosis in the condition vector, as a means to add more specificity to the type of output that is generated. Our generative framework illustrates a potential application for generating synthetic training data that would be beneficial for improving deep learning frameworks that benefit from increased dataset sizes. Significant volume and surface area changes w.r.t. to AD diagnosis were identified, particularly in the amygdala, caudate, nucleus accumbens, right putamen, thalamus, and most importantly the hippocampus, an area of the brain highly correlated with AD. Our prior work using spectral filters (Azcona et al., 2020) utilizes the same subcortical structures, in addition to the cortex, to perform the same AD classification task. However, during our analysis, we observed frequent GPU memory issues with training a CVAE model using the cortical surface. A major degradation in the output quality of reconstructed/generated cortical surfaces with our generative framework was also observed. As Gutiérrez-Becker et al. (2021) point out, modeling a structure with a more complex geometry, e.g., the cortex, requires a larger number of points that may lead to GPU memory constraints. Additionally, the gyrification of the cortical surface is much more complex and may require additional methods that generalize better to 3D mesh structures with complex sulci and gyri.

## 5. Conclusions and Future Work

To the best of our knowledge, no existing works have investigated brain shape in regards to AD pathology using discriminative *and* generative networks that learn and operate *directly on surface meshes* by way of geometric deep learning. Our framework is constructed by a variety of modular computational blocks that are used by both our discriminative and generative models. Notably, our convolutional encoder learns effective shape descriptors that can be used for AD classification by our discriminative model. Our first analysis demonstrates an improvement in AD classification performance using the *same model* with varying input types: (a) single subcortical region, (b) subcortical regions within a single hemisphere, and (c) bilateral subcortical regions. Our results demonstrate a clear advantage to the joint modeling of multiple subcortical structures for *in-vivo* AD classification.

Our discriminative model also outperforms alternative shape descriptor methods in our baseline comparison. Additionally, our adaptation of Grad-CAM to 3D meshes provides context as to which subcortical brain regions are driving our AD classification results. Our class activation maps (CAMs) are in accordance with the literature on morphological changes observed in the brains of subjects with AD. Our CAMs make our classification results more transparent by producing visual explanations. Improving clinical confidence and reliability in automated discriminative methods, can be approached by contextualizing a model’s reasoning about its beliefs and actions for clinicians to trust and use.

Additionally, our generative model’s decoder module is able to reconstruct 3D mesh inputs from their low-dimensional shape descriptors obtained by the encoder. More importantly, in using a variational approach, we’re able to learn a probabilistic latent space that can be sampled from to generate synthetic samples for each subcortical structure w.r.t. phenotype information, in particular: AD diagnosis. The endemic nature of medical imaging data, particularly within neuroimaging, attributes to scarcity of open-access neuroimaging databases. Our generative model is able to generate realistic-looking synthetic examples, which may be used to train other deep learning approaches that often require large datasets and annotated data is limited.

Our proposed discriminative network can be further tailored to fuse other phenotypic data for AD classification; including but not limited to: chronological age, sex assignment at birth, genotype data, etc. Phenotype features can also be used as additional conditional priors in our generative framework, adding additional constraints for synthesizing personalized samples. Natural extensions of this work could include (1) expanding the classification task to sub-typing different stages of mild cognitive impairment (early versus late), (2) using spiral convolution within a recurrent neural network framework for longitudinal predictions related to AD, and (3) experimenting with generating template-registered 3D meshes from MRI volume inputs using a spiral convolutional decoder framework to automate the mesh extraction preprocessing steps.

## CRediT Authorship Contribution Statement

**Emanuel A. Azcona:** Conceptualization, Methodology, Software, Validation, Visualization, Investigation, Writing original draft, review, & editing. **Pierre Besson:** Conceptualization, Software, Validation, Investigation, Data Curation, Resources, Supervision, Writing - review & editing. **Yunan Wu:** Software, Validation, Investigation, Writing - review & editing. **Ajay S. Kurani:** Methodology, Validation, Resources, Writing - review & editing. **S. Kathleen Bandt:** Conceptualization, Methodology, Validation, Resources, Writing - review & editing. **Todd B. Parrish:** Validation, Resources, Supervision, Funding acquisition, Writing - review & editing. **Aggelos K. Katsaggelos:** Validation, Resources, Supervision, Funding acquisition, Writing - review & editing.

## Acknowledgements

This work was funded in part by the Biomedical Data Driven Discovery Training Grant from the National Library of Medicine (5T32LM012203) through Northwestern University, and the National Institute on Aging. The authors would also like to thank the QUEST High Performance Computing Cluster at Northwestern University for computational resources.

Data collection and sharing for this project was funded by the Alzheimer’s Disease Neuroimaging Initiative (ADNI) (National Institutes of Health Grant U01 AG024904) and DOD ADNI (Department of Defense award number W81XWH-12-2-0012). ADNI is funded by the National Institute on Aging, the National Institute of Biomedical Imaging and Bioengineering, and through generous contributions from the following: AbbVie, Alzheimer’s Association; Alzheimer’s Drug Discovery Foundation; Araclon Biotech; BioClinica, Inc.; Biogen; Bristol-Myers Squibb Company; CereSpir, Inc.; Cogstate; Eisai Inc.; Elan Pharmaceuticals, Inc.; Eli Lilly and Company; EuroImmun; F. Hoffman-La Roche Ltd and its affiliated company Genentech, Inc.; Fujirebio; GE Healthcare; IXICO Ltd; Janssen Alzheimer Immunotherapy Research & Development, LLC.; Johnson & Johnson Pharmaceutical Research & Development LLC.; Lumosity; Lundbeck; Merck & Co., Inc.; Meso Scale Diagnostics, LLC.; NeuroRx Research; Neurotrack Technologies; Novartis Pharmaceuticals Corporation; Pfizer Inc.; Piramal Imaging; Servier; Takeda Pharmaceutical Company; and Transition Therapeutics. The Canadian Institutes of Health Research is providing funds to support ADNI clinical sites in Canada. Private sector contributions are facilitated by the Foundation for the National Institutes of Health (www.fnih.org). The grantee organization is the Northern California Institute for Research and Education, and the study is coordinated by the Alzheimer’s Therapeutic Research Institute at the University of Southern California. ADNI data are disseminated by the Laboratory for Neuro Imaging at the University of Southern California.

## References

Achlioptas, P., Diamanti, O., Mitliagkas, I., Guibas, L., 2018. Learning representations and generative models for 3D point clouds, in: Dy, J., Krause, A. (Eds.), Proceedings of the 35th International Conference on Machine Learning, PMLR. pp. 40–49.

Azcona, E.A., Besson, P., Wu, Y., Punjabi, A., Martersteck, A., Dravid, A., Parrish, T.B., Bandt, S.K., Katsaggelos, A.K., 2020. Interpretation of brain morphology in association to alzheimer’s disease dementia classification using graph convolutional networks on triangulated meshes, in: International Workshop on Shape in Medical Imaging, Springer. pp. 95–107.

Barber, R., McKeith, I., Ballard, C., O’Brien, J., 2002. Volumetric mri study of the caudate nucleus in patients with dementia with lewy bodies, alzheimer’s disease, and vascular dementia. Journal of Neurology, Neurosurgery & Psychiatry 72, 406–407.

Bessadok, A., Mahjoub, M.A., Rekik, I., 2019. Hierarchical adversarial connectomic domain alignment for target brain graph prediction and classification from a source graph, in: International Workshop on PRedictive Intelligence In MEdicine, Springer. pp. 105–114.

Bessadok, A., Mahjoub, M.A., Rekik, I., 2021. Brain graph synthesis by dual adversarial domain alignment and target graph prediction from a source graph. Medical Image Analysis 68, 101902.

Bessadok, A., Rekik, I., 2018. Intact connectional morphometricity learning using multi-view morphological brain networks with application to autism spectrum disorder, in: International Workshop on Connectomics in Neuroimaging, Springer. pp. 38–46.

Besson, P., Lopes, R., Leclerc, X., Derambure, P., Tyvaert, L., 2014. Intra-subject reliability of the high-resolution whole-brain structural connectome. NeuroImage 102, 283–293.

Besson, P., Parrish, T., Katsaggelos, A.K., Bandt, S.K., 2020. Geometric deep learning on brain shape predicts sex and age. bioRxiv doi:10.1101/2020.06.29.177543.

Bouritsas, G., Bokhnyak, S., Ploumpis, S., Bronstein, M., Zafeiriou, S., 2019. Neural 3d morphable models: Spiral convolutional networks for 3d shape representation learning and generation, in: Proceedings of the IEEE/CVF International Conference on Computer Vision, pp. 7213–7222.

Braak, H., Braak, E., 1991. Neuropathological stageing of alzheimer-related changes. Acta neuropathologica 82, 239–259.

Brignell, C.J., Dryden, I.L., Gattone, S.A., Park, B., Leask, S., Browne, W.J., Flynn, S., 2010. Surface shape analysis with an application to brain surface asymmetry in schizophrenia. Biostatistics 11, 609–630.

Bronstein, M.M., Bruna, J., LeCun, Y., Szlam, A., Vandergheynst, P., 2017. Geometric deep learning: going beyond euclidean data. IEEE Signal Processing Magazine 34, 18–42.

Choi, H., Kang, H., Lee, D.S., Initiative, A.D.N., et al., 2018. Predicting aging of brain metabolic topography using variational autoencoder. Frontiers in aging neuroscience 10, 212.

Clevert, D.A., Unterthiner, T., Hochreiter, S., 2015. Fast and accurate deep network learning by exponential linear units (elus). International Conference on Learning Representations (ICLR).

Corballis, M.C., 2014. Left brain, right brain: facts and fantasies. PLoS Biol 12, e1001767.

Corso, G., Cavalleri, L., Beaini, D., Lió, P., Velic?ković, P., 2020. Principal neighbourhood aggregation for graph nets. arXiv preprint 2004.05718

Dale, A.M., Fischl, B., Sereno, M.I., 1999. Cortical surface-based analysis: I. segmentation and surface reconstruction. Neuroimage 9, 179–194.

Dale, A.M., Sereno, M.I., 1993. Improved localizadon of cortical activity by combining eeg and meg with mri cortical surface reconstruction: a linear approach. Journal of cognitive neuroscience 5, 162–176.

Dawson-Haggerty, et al., 2019. trimesh. URL: https://trimsh.org/.

Defferrard, M., Bresson, X., Vandergheynst, P., 2016. Convolutional neural networks on graphs with fast localized spectral filtering, in: Proceedings of the 30th International Conference on Neural Information Processing Systems, pp. 3844–3852.

Deng, J., Dong, W., Socher, R., Li, L.J., Li, K., Fei-Fei, L., 2009. Imagenet: A large-scale hierarchical image database, in: 2009 IEEE conference on computer vision and pattern recognition, Ieee. pp. 248–255.

Derflinger, S., Sorg, C., Gaser, C., Myers, N., Arsic, M., Kurz, A., Zimmer, C., Wohlschläger, A., Mühlau, M., 2011. Grey-matter atrophy in alzheimer’s disease is asymmetric but not lateralized. Journal of Alzheimer’s Disease 25, 347–357.

Dickerson, B.C., Goncharova, I., Sullivan, M., Forchetti, C., Wilson, R., Bennett, D., Beckett, L.A., deToledo Morrell, L., 2001. Mri-derived entorhinal and hippocampal atrophy in incipient and very mild alzheimer’s disease. Neurobiology of aging 22, 747–754.

Dubois, B., Feldman, H.H., Jacova, C., DeKosky, S.T., Barberger-Gateau, P., Cummings, J., Delacourte, A., Galasko, D., Gauthier, S., Jicha, G., et al., 2007. Research criteria for the diagnosis of alzheimer’s disease: revising the nincds–adrda criteria. The Lancet Neurology 6, 734–746.

Ferrarini, L., Palm, W.M., Olofsen, H., van Buchem, M.A., Reiber, J.H., Admiraal-Behloul, F., 2006. Shape differences of the brain ventricles in alzheimer’s disease. Neuroimage 32, 1060–1069.

Fey, M., Lenssen, J.E., 2019. Fast graph representation learning with pytorch geometric. arXiv preprint 1903.02428.

Fischl, B., 2012. Freesurfer. Neuroimage 62, 774–781.

Fischl, B., Salat, D.H., Busa, E., Albert, M., Dieterich, M., Haselgrove, C., Van Der Kouwe, A., Killiany, R., Kennedy, D., Klaveness, S., et al., 2002. Whole brain segmentation: automated labeling of neuroanatomical structures in the human brain. Neuron 33, 341–355.

Fischl, B., Sereno, M.I., Dale, A.M., 1999a. Cortical surface-based analysis: Ii: inflation, flattening, and a surface-based coordinate system. Neuroimage 9, 195–207.

Fischl, B., Sereno, M.I., Tootell, R.B., Dale, A.M., 1999b. High-resolution intersubject averaging and a coordinate system for the cortical surface. Human brain mapping 8, 272–284.

Fornito, A., Zalesky, A., Breakspear, M., 2015. The connectomics of brain disorders. Nature Reviews Neuroscience 16, 159–172.

Frings, L., Hellwig, S., Spehl, T.S., Bormann, T., Buchert, R., Vach, W., Minkova, L., Heimbach, B., Klöppel, S., Meyer, P.T., 2015. Asymmetries of amyloid-β burden and neuronal dysfunction are positively correlated in alzheimer’s disease. Brain 138, 3089–3099.

Frisoni, G.B., Fox, N.C., Jack, C.R., Scheltens, P., Thompson, P.M., 2010. The clinical use of structural mri in alzheimer disease. Nature Reviews Neurology 6, 67–77.

Garland, M., Heckbert, P.S., 1997. Surface simplification using quadric error metrics, in: Proceedings of the 24th annual conference on Computer graphics and interactive techniques, pp. 209–216.

Gilmer, J., Schoenholz, S.S., Riley, P.F., Vinyals, O., Dahl, G.E., 2017. Neural message passing for quantum chemistry, in: International Conference on Machine Learning, PMLR. pp. 1263–1272.

Göktaş, A.S., Bessadok, A., Rekik, I., 2020. Residual embedding similarity-based network selection for predicting brain network evolution trajectory from a single observation, in: International Workshop on PRedictive Intelligence In MEdicine, Springer. pp. 12–23.

Gong, S., Chen, L., Bronstein, M., Zafeiriou, S., 2019. Spiralnet++: A fast and highly efficient mesh convolution operator, in: Proceedings of the IEEE/CVF International Conference on Computer Vision Workshops, pp. 0–0.

Goodfellow, I.J., Pouget-Abadie, J., Mirza, M., Xu, B., Warde-Farley, D., Ozair, S., Courville, A., Bengio, Y., 2014. Generative adversarial nets, in: Proceedings of the 27th International Conference on Neural Information Processing Systems - Volume 2, MIT Press. p. 2672–2680.

Gurbuz, M.B., Rekik, I., 2020. Deep graph normalizer: A geometric deep learning approach for estimating connectional brain templates, in: International Conference on Medical Image Computing and Computer-Assisted Intervention, Springer. pp. 155–165.

Gutiérrez-Becker, B., Sarasua, I., Wachinger, C., 2021. Discriminative and generative models for anatomical shape analysis on point clouds with deep neural networks. Medical Image Analysis 67, 101852.

He, K., Zhang, X., Ren, S., Sun, J., 2016. Deep residual learning for image recognition, in: Proceedings of the IEEE conference on computer vision and pattern recognition, pp. 770–778.

Ioffe, S., Szegedy, C., 2015. Batch normalization: Accelerating deep network training by reducing internal covariate shift, in: Proceedings of the 32nd International Conference on Machine Learning, PMLR, Lille, France. pp. 448–456.

de Jong, L.W., van der Hiele, K., Veer, I.M., Houwing, J., Westendorp, R., Bollen, E., de Bruin, P.W., Middelkoop, H., van Buchem, M.A., van der Grond, J., 2008. Strongly reduced volumes of putamen and thalamus in alzheimer’s disease: an mri study. Brain 131, 3277–3285.

Joyce, J.M., 2011. Kullback-leibler divergence. International Encyclopedia of Statistical Science, 720–722.

Keilp, J.G., Alexander, G.E., Stern, Y., Prohovnik, I., 1996. Inferior parietal perfusion, lateralization, and neuropsychological dysfunction in alzheimer’s disease. Brain and cognition 32, 365–383.

Kim, H., Mansi, T., Bernasconi, N., 2013. Disentangling hippocampal shape anomalies in epilepsy. Frontiers in neurology 4, 131.

Kingma, D.P., Welling, M., 2014. Auto-encoding variational bayes. 2nd International Conference on Learning Representations, (ICLR) 2014, Banff, AB, Canada, April 14-16, 2014, Conference Track Proceedings.

Kipf, T.N., Welling, M., 2017. Semi-supervised classification with graph convolutional networks. International Conference on Learning Representations (ICLR).

Klein-Koerkamp, Y.A Heckemann, R. T Ramdeen, K., Moreaud, O., Keignart, S., Krainik, A., Hammers, A., Baciu, M., Hot, P., disease Neuroimaging Initiative, A., et al., 2014. Amygdalar atrophy in early alzheimer’s disease. Current Alzheimer Research 11, 239–252.

Klöppel, S., Stonnington, C.M., Chu, C., Draganski, B., Scahill, R.I., Rohrer, J.D., Fox, N.C., Jack Jr, C.R., Ashburner, J., Frackowiak, R.S., 2008. Automatic classification of mr scans in alzheimer’s disease. Brain 131, 681–689.

Kolotouros, N., Pavlakos, G., Daniilidis, K., 2019. Convolutional mesh regression for single-image human shape reconstruction, in: Proceedings of the IEEE/CVF Conference on Computer Vision and Pattern Recognition, pp. 4501–4510.

LeCun, Y., Boser, B., Denker, J.S., Henderson, D., Howard, R.E., Hubbard, W., Jackel, L.D., 1989. Backpropagation applied to handwritten zip code recognition. Neural computation 1, 541–551.

Ledig, C., Schuh, A., Guerrero, R., Heckemann, R.A., Rueckert, D., 2018. Structural brain imaging in alzheimer’s disease and mild cognitive impairment: biomarker analysis and shared morphometry database. Scientific reports 8, 1–16.

Lim, I., Dielen, A., Campen, M., Kobbelt, L., 2018. A simple approach to intrinsic correspondence learning on unstructured 3d meshes, in: Proceedings of the European Conference on Computer Vision (ECCV) Workshops, pp. 0–0.

Litany, O., Bronstein, A., Bronstein, M., Makadia, A., 2018. Deformable shape completion with graph convolutional autoencoders, in: Proceedings of the IEEE conference on computer vision and pattern recognition, pp. 1886– 1895.

Long, X., Zhang, L., Liao, W., Jiang, C., Qiu, B., Initiative, A.D.N., 2013. Distinct laterality alterations distinguish mild cognitive impairment and alzheimer’s disease from healthy aging: Statistical parametric mapping with high resolution mri. Human brain mapping 34, 3400–3410.

Loshchilov, I., Hutter, F., 2017. Decoupled weight decay regularization. arXiv preprint 1711.05101.

Lu, L., Shin, Y., Su, Y., Karniadakis, G.E., 2020. Dying relu and initialization: Theory and numerical examples. Communications in Computational Physics 28, 1671–1706.

Mann, D., 1991. The topographic distribution of brain atrophy in alzheimer’s disease. Acta neuropathologica 83, 81–86.

Marton, Z.C., Rusu, R.B., Beetz, M., 2009. On fast surface reconstruction methods for large and noisy point clouds, in: 2009 IEEE international conference on robotics and automation, IEEE. pp. 3218–3223.

McKight, P.E., Najab, J., 2010. Kruskal-wallis test. The Corsini Encyclopedia of Psychology.

Nebli, A., Rekik, I., 2020. Gender differences in cortical morphological networks. Brain imaging and behavior 14, 1831–1839.

Nelson, P.T., Head, E., Schmitt, F.A., Davis, P.R., Neltner, J.H., Jicha, G.A., Abner, E.L., Smith, C.D., Van Eldik, L.J., Kryscio, R.J., et al., 2011. Alzheimer’s disease is not “brain aging”: neuropathological, genetic, and epidemiological human studies. Acta neuropathologica 121, 571–587.

Ng, B., Toews, M., Durrleman, S., Shi, Y., 2014. Shape analysis for brain structures. Shape Analysis in Medical Image Analysis, 3–49.

Niethammer, M., Reuter, M., Wolter, F.E., Bouix, S., Peinecke, N., Koo, M.S., Shenton, M.E., 2007. Global medical shape analysis using the laplace-beltrami spectrum, in: International Conference on Medical Image Computing and Computer-Assisted Intervention, Springer. pp. 850–857.

Petersen, R.C., Aisen, P.S., Beckett, L.A., Donohue, M.C., Gamst, A.C., Harvey, D.J., Jack, C.R., Jagust, W.J., Shaw, L.M., Toga, A.W., Trojanowski, J.Q., Weiner, M.W., 2010. Alzheimer’s disease neuroimaging initiative (adni). Neurology 74, 201–209. doi:10.1212/WNL.0b013e3181cb3e25.

Punjabi, A., Martersteck, A., Wang, Y., Parrish, T.B., Katsaggelos, A.K., Initiative, A.D.N., 2019. Neuroimaging modality fusion in alzheimer’s classification using convolutional neural networks. PloS one 14, e0225759.

Qi, C.R., Su, H., Mo, K., Guibas, L.J., 2017a. Pointnet: Deep learning on point sets for 3d classification and segmentation, in: Proceedings of the IEEE conference on computer vision and pattern recognition, pp. 652–660.

Qi, C.R., Yi, L., Su, H., Guibas, L.J., 2017b. Pointnet++ deep hierarchical feature learning on point sets in a metric space, in: Proceedings of the 31st International Conference on Neural Information Processing Systems, pp. 5105–5114.

Ranjan, A., Bolkart, T., Sanyal, S., Black, M.J., 2018. Generating 3d faces using convolutional mesh autoencoders, in: Proceedings of the European Conference on Computer Vision (ECCV), pp. 704–720.

Reuter, M., Wolter, F.E., Peinecke, N., 2006. Laplace–beltrami spectra as ‘shape-dna’of surfaces and solids. Computer-Aided Design 38, 342–366.

Selvaraju, R.R., Cogswell, M., Das, A., Vedantam, R., Parikh, D., Batra, D., 2017. Grad-cam: Visual explanations from deep networks via gradient-based localization, in: Proceedings of the IEEE international conference on computer vision, pp. 618–626.

Shakeri, M., Lombaert, H., Datta, A.N., Oser, N., Létourneau-Guillon, L., Lapointe, L.V., Martin, F., Malfait, D., Tucholka, A., Lippé, S., et al., 2016. Statistical shape analysis of subcortical structures using spectral matching. Computerized Medical Imaging and Graphics 52, 58–71.

Sohn, K., Lee, H., Yan, X., 2015. Learning structured output representation using deep conditional generative models. Advances in neural information processing systems 28, 3483–3491.

Sserwadda, A., Rekik, I., 2021. Topology-guided cyclic brain connectivity generation using geometric deep learning. Journal of Neuroscience Methods 353, 108988.

Szegedy, C., Vanhoucke, V., Ioffe, S., Shlens, J., Wojna, Z., 2016. Rethinking the inception architecture for computer vision, in: Proceedings of the IEEE conference on computer vision and pattern recognition, pp. 2818–2826.

TaubinÝ, G., 2000. Geometric signal processing on polygonal meshes. Proceedings of EUROGRAPHICS 2000: state of the art report.

Thompson, P.M., Hayashi, K.M., Dutton, R.A., Chiang, M.C., Leow, A.D., Sowell, E.R., De Zubicaray, G., Becker, J.T., Lopez, O.L., Aizenstein, H.J., et al., 2007. Tracking alzheimer’s disease. Annals of the New York Academy of Sciences 1097, 183.

Tondelli, M., Wilcock, G.K., Nichelli, P., De Jager, C.A., Jenkinson, M., Zamboni, G., 2012. Structural mri changes detectable up to ten years before clinical alzheimer’s disease. Neurobiology of aging 33, 825–e25.

Van Rossum, G., Drake Jr, F.L., 1995. Python tutorial. Centrum voor Wiskunde en Informatica Amsterdam, The Netherlands.

Wachinger, C., Golland, P., Kremen, W., Fischl, B., Reuter, M., Initiative, A.D.N., et al., 2015. Brainprint: A discriminative characterization of brain morphology. NeuroImage 109, 232–248.

Wang, N., Zhang, Y., Li, Z., Fu, Y., Liu, W., Jiang, Y.G., 2018. Pixel2mesh: Generating 3d mesh models from single rgb images, in: Proceedings of the European Conference on Computer Vision (ECCV), pp. 52–67.

Wickramasinghe, U., Remelli, E., Knott, G., Fua, P., 2020. Voxel2mesh: 3d mesh model generation from volumetric data, in: International Conference on Medical Image Computing and Computer-Assisted Intervention, Springer. pp. 299–308.

Wu, J., Zhang, C., Xue, T., Freeman, W.T., Tenenbaum, J.B., 2016. Learning a probabilistic latent space of object shapes via 3d generative-adversarial modeling, in: Proceedings of the 30th International Conference on Neural Information Processing Systems, Curran Associates Inc., Red Hook, NY, USA. p. 82–90.

Wu, Y., Besson, P., Azcona, E.A., Bandt, S.K., Parrish, T.B., Breiter, H.C., Katsaggelos, A.K., 2020a. Novel age-dependent cortico-subcortical morhologic interactions predict fluid intelligence: A multi-cohort geometric deep learning study. bioRxiv.

Wu, Z., Pan, S., Chen, F., Long, G., Zhang, C., Philip, S.Y., 2020b. A comprehensive survey on graph neural networks. IEEE transactions on neural networks and learning systems.

Xie, Y., Li, S., Yang, C., Wong, R.C.W., Han, J., 2020. When do gnns work: Understanding and improving neighborhood aggregation. IJCAI’20: Proceedings of the Twenty-Ninth International Joint Conference on Artificial Intelligence, {IJCAI} 2020 2020.

Yang, J., Zhu, Q., Zhang, R., Huang, J., Zhang, D., 2020. Unified brain network with functional and structural data, in: International Conference on Medical Image Computing and Computer-Assisted Intervention, Springer. pp. 114– 123.

Yu, F., Koltun, V., 2016. Multi-scale context aggregation by dilated convolutions. International Conference on Learning Representations (ICLR).

Zhang, L., Wang, L., Zhu, D., 2020. Recovering brain structural connectivity from functional connectivity via multi-gcn based generative adversarial network, in: International Conference on Medical Image Computing and Computer-Assisted Intervention, Springer. pp. 53–61.

